# Multiple neuronal networks coordinate *Hydra* mechanosensory behavior

**DOI:** 10.1101/2020.10.16.343384

**Authors:** Krishna N. Badhiwala, Abby S. Primack, Celina E. Juliano, Jacob T. Robinson

## Abstract

*Hydra vulgaris* is an emerging model organism for neuroscience due to its small size, transparency, genetic tractability, and regenerative nervous system; however, fundamental properties of its sensorimotor behaviors remain unknown. Here, we use microfluidic devices combined with fluorescent calcium imaging and surgical resectioning to study how the diffuse nervous system coordinates *Hydra*’s mechanosensory response. Mechanical stimuli cause animals to contract, and we find this response relies on at least two distinct networks of neurons in the oral and aboral regions of the animal. Different activity patterns arise in these networks depending on whether the animal is contracting spontaneously or contracting in response to mechanical stimulation. Together, these findings improve our understanding of how *Hydra*’s diffuse nervous system coordinates sensorimotor behaviors. These insights help reveal how sensory information is processed in an animal with a diffuse, radially symmetric neural architecture unlike the dense, bilaterally symmetric nervous systems found in most model organisms.

## Introduction

Discovering the fundamental principles of neural activity and behaviors requires studying the nervous systems of diverse organisms. Animals have evolved different neural structures like the nerve net (e.g. *Hydra*), nerve cords and ganglia (e.g. *C. elegans*, *Aplysia*, planaria), and brain (e.g. *Drosophila*, zebrafish, rodents, and primates). Despite the vastly different structures, many behaviors are conserved across species, including sensorimotor responses^1–6^ and sleep^7–15^. By comparing neural circuits that support similar behaviors despite different architectures, we can discover organizational principles of neural circuits that reflect millions of years of evolutionary pressure. While there are many potential organisms that would support this type of comparative neuroscience, only a small group of animals have the qualities to support laboratory experiments: short generation span, ease of breeding and manipulation in laboratory conditions, small and compact size, optical transparency, and a well-developed genetic toolkit with a complete spatial and molecular map of the nervous system.

Transparent, millimeter-sized animals in particular offer a number of advantages for neuroscientists because it is possible to image neural activity throughout the entire nervous system using genetically-encoded calcium or voltage-sensitive fluorescent proteins^16–25^. In addition, some millimeter-sized animals are compatible with microfluidic devices for precise environmental control and microscopy techniques that offer cellular-resolution functional imaging of the entire nervous system. These properties, combined with genetic tractability, provide a powerful way of revealing neuronal dynamics across the entire nervous system (not just a small region) during behaviors. For instance, whole nervous system imaging of confined or freely moving animals has revealed the neuronal dynamics underlying locomotion^19, 26^ and sensory motivated global state transitions in *C. elegans*^27, 28^, responses to noxious odor and visuomotor behaviors in zebrafish^19–24^, responses to light and odor in *Drosophila*^18, 29^, and neuronal ensembles correlated with basal behaviors and response to light in *Hydra*^30, 31^.

*Hydra* is unique among the small, transparent organisms discussed above due to its regenerative ability and highly dynamic nervous system. While most small, transparent model systems (like *C. elegans* or zebrafish larvae) suffer permanent behavioral deficits from the loss of one or a few neurons^32–36^, *Hydra* can completely recover from a significant neuronal loss to regain normal contractile behavior in as little as ∼ 48 hours^37–39^. This radially symmetric freshwater cnidarian has a nervous system composed of two diffuse networks of neurons, one embedded in the endoderm and another embedded in the ectoderm^40, 41^. While *Hydra’s* diffuse nerve net is highly dynamic with continuous cellular turnover and migration^42, 43^, regions with increased neuron density resembling nerve rings have a comparatively lower neuronal turnover (Figure 1a)^44–47^. One of these regions is in the oral end in the apex above the ring of tentacles (“hypostomal nerve ring”), and another is in the aboral end in the foot (“peduncle nerve ring”) (Figure 1a). Recent single-cell RNA sequencing has provided a complete molecular and spatial map of the *Hydra* nervous system, including identification of unique cell-type specific biomarkers to generate new transgenic models^48^. This existing molecular and spatial map of the nervous system suggests there is no overlap in the neuronal cell types that make up the hypostomal and peduncle nerve rings, and the distribution of the cell types varies along the length of the body (Figure 1b). Finally, the demonstration of microfluidic and transgenic tools combined with *Hydra’s* dynamic yet ‘simple’ neural architecture has enabled observations of basal and sensory motivated behaviors in the regenerating nervous system^31^.

**Figure 1:**
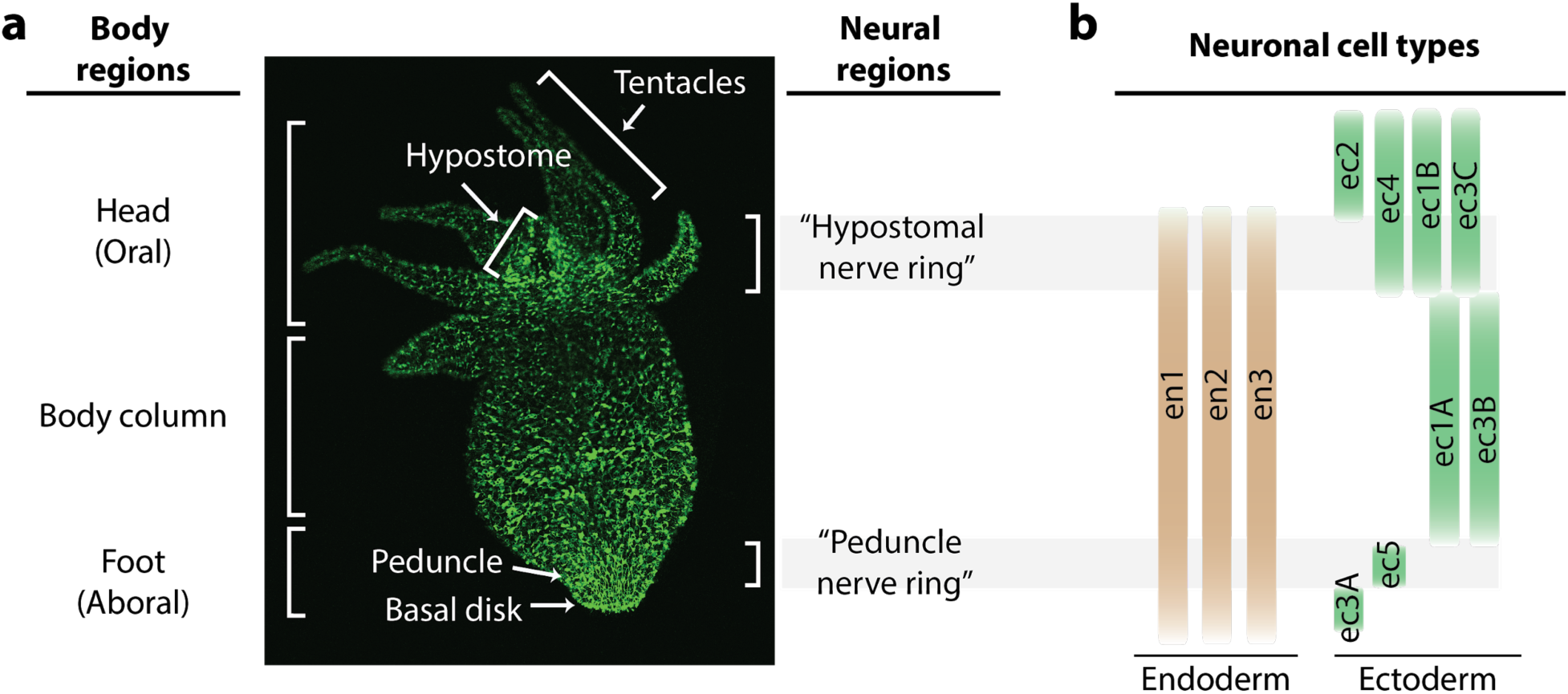
Distribution of neurons in the *Hydra* nerve net. a) Fluorescent image of *Hydra* nervous system. Green fluorescent protein (GFP) is expressed in neurons and neuronal progenitors (nGreen transgenic line^48^). Body anatomy is annotated on the left. White arrows indicate the body parts: hypostome, tentacles, peduncle, and basal disk. High neuronal density regions are annotated on the right. b) Distribution of neuronal cell types varying longitudinally along the body, with endodermal nerve net cell types in tan and ectodermal nerve net cell types in green. Cell types were identified through single cell RNA sequencing^48^.

To better establish *Hydra* as a model organism for comparative neuroscience, it is critical to understand their basic sensorimotor behaviors, such as response to touch. While it is well documented that *Hydra* contract when mechanically agitated or poked with a pipette^49–53^, we found no quantitative reports of how this behavior depends on stimulus intensity or is mediated by neural activity. Although significant insights in *Hydra* behavior have been made over the last several decades with simple methodologies and manual observations, including that the tentacles and/or the hypostome are needed for mechanosensory response, these experiments lack quantitative characterization of neuronal or behavioral response. Forceps-induced touch allows local stimulation, but it is difficult to control the force applied manually^54, 55^. While stimulation with mechanical agitation allows control over stimulus intensity, these observations are limited to changes in body lengths^52^.

Here, we use whole animal functional imaging combined with resection studies to discover that despite the apparently diffuse nerve net in *Hydra*, these animals process sensorimotor responses in specialized regional networks. To study the mechanosensory response, we first developed a microfluidic system to apply a local mechanical stimulus and quantify *Hydra’s* behavioral and neural response. We then measured these responses in the absence of select regions of the body and found at least one of the neuron-rich regions, the hypostome (oral) or the peduncle (aboral), is required to coordinate spontaneous contractions, though the oral network plays a more significant role. We found a significant reduction in the mechanosensory response with the removal of the hypostome, the region where sensory information is likely processed. These sensorimotor experiments combined with whole-animal neural and epitheliomuscular imaging reveal that *Hydra* is capable of receiving sensory information along the body column; however, the oral region is necessary for coordinating the motor response.

## Results

### *Hydra’s* mechanosensory response is dependent on stimulus intensity

To better understand sensory information processing in *Hydra*, we developed a double-layer microfluidic system that can apply a local mechanical stimulus while we image the response of the entire nervous system using fluorescence microscopy (Figure 2, Video 1). This local mechanical stimulation is made possible by push-down microfluidic valves (Figure 2a) that deliver mechanical stimuli to a portion of the *Hydra* body with precise temporal and spatial control (see “Methods”). For all experiments, we pressurized a valve (400 µm diameter) that was directly above the animal (for 1 sec every 31 sec, see “Methods”) to stimulate the body column while simultaneously performing functional calcium imaging (Figure 2b). We selected the middle of the body for stimulation region to help ensure that we stimulated roughly the same region of the animal throughout each experiment. This choice was based on the observation that the body column region was relatively stationary, whereas the oral and aboral extremities had large displacements during body contractions and elongations.

**Figure 2:**
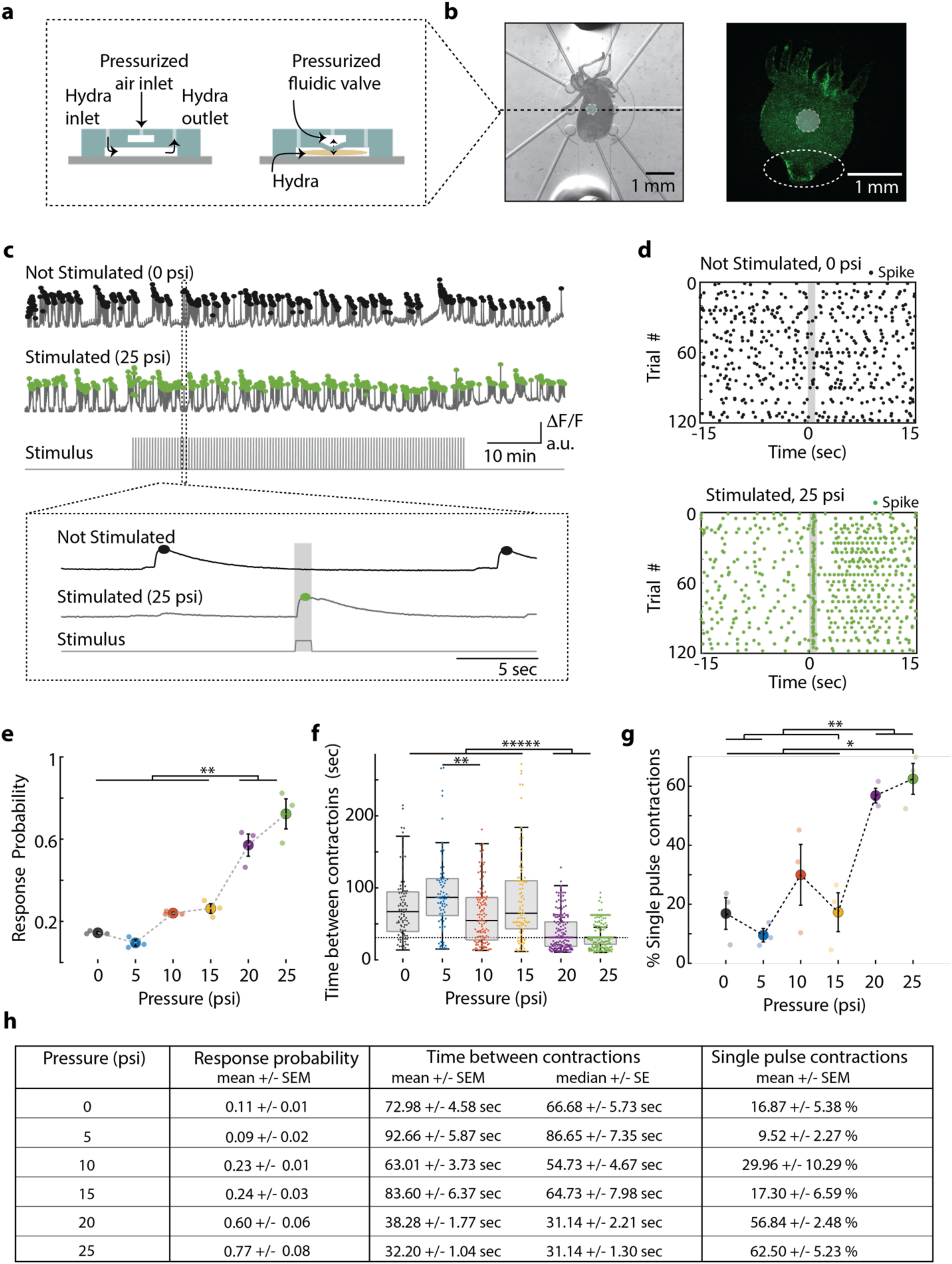
*Hydra’s* neuronal response depends on the mechanical stimulus intensity. a) (left) Side view of double-layer microfluidic device for mechanical stimulation. (right) Device with pressurized valve. *Hydra* is immobilized in the bottom *Hydra* layer, and pressurized air supplied into the valve layer causes the circular membrane (400 µm diameter) to push down on *Hydra*. b) (left) Brightfield image of *Hydra* immobilized in the bottom layer of the chip and the arrangement of micro-valves on the top layer. Micro-valve used for stimulation falsely colored with a light blue circle. (right) Fluorescent image of *Hydra* with pan-neuronal expression of GCaMP6s. White dashed circle marks the peduncle region of interest (ROI) used for quantifying calcium fluorescence changes. c-d) Representative calcium fluorescence activity in the peduncle region from an animal not stimulated and animal mechanically stimulated with 25 psi. Black and green dots indicate fluorescence (calcium) spikes. Gray shaded regions indicate stimulus ‘on’ time (also “response window”). c) Fluorescence (calcium) trace from *Hydra* not stimulated (top) and stimulated with 25 psi (middle). Stimulation protocol in gray (bottom trace): 20 min no stimulation, 1 hr of repeated stimulation (1 sec ‘on’, 30 sec ‘off’) and 20 min no stimulation. Stimulus ‘on’ times indicated with vertical lines. Magnification of 30 sec fluorescence and stimulation protocol trace from one stimulation trial. d) Raster plot of stimulus time aligned spiking activity from multiple trials superimposed for *Hydra* not stimulated (top) and stimulated with 25 psi (bottom). e). Mechanosensory response probability, fraction of trials (out of 119 total) that have at least one calcium spike (also contraction pulse) occurring during the 1 sec response window (gray shaded region) when valve is pressurized. Large circles indicate average probability from all animals (N=3) combined for each condition. Small circles indicate probability from a single animal. Significant pairwise comparisons are shown with brackets (one-way ANOVA with post-hoc Bonferroni correction). f) Time interval between body contractions under each condition. Dashed line represents the time interval between stimuli (∼31 sec). Brackets indicate significant differences in a Kruskal-Wallis test with post-hoc Dunn-Sidak correction. g) Percent of all body contractions that are a single pulse; brackets show significant pairwise comparisons from a one-way ANOVA with post-hoc Bonferroni correction. Error bars are standard error of mean (SEM); N = 3 *Hydra* for each condition; * = p < 0.05, ** = p < 0.01, ***** = p < 0.00001. h) Table summarizing the mechanosensory response probability, time between contractions, and percent of contractions that are single pulses for each stimulus intensity (mean ± SEM or median ± SE). Source data for the quantitative characterization of mechanosensory response are available in the Figure 2 – source data 1.

Experiments showed that this stimulation paradigm delivered a local mechanical stimulation with most of the mechanical force localized to a radius of approximately 250 µm around the microfluidic valve. To measure the locality of this stimulus we performed an experiment using transgenic *Hydra* (nGreen)^48^ expressing GFP pan-neuronally (and in neural progenitors) and tracked the position and fluorescence intensity (GFP) from individual neurons during mechanical stimulation (N = 222 neurons over 1 min). When the microfluidic valve was pressurized to deliver mechanical stimulation, we found significantly increased average cellular (or tissue) displacement (p < 0.001). Further analysis of the cellular movements showed the spatial distribution of mechanical force from the stimulation was primarily experienced by the neurons directly under the valve (Figure 2 – Figure Supplement 1). Specifically, we found that the tissue directly below the valve was compressed (z direction) when mechanically stimulated - the neurons directly under the valve had a small magnitude of lateral (x-y direction) displacement. The tissue bordering the valve was stretched away from the valve center - the neurons in the neighboring regions around the valve had the largest lateral displacement. This lateral displacement decreased for neurons that were farther from the center of the valve. Neurons more than 750 µm from the microfluidic valve center showed a negligible displacement of less than 5 µm (95% CI lower bound = 5.8 µm), which is ∼550% less than the displacement of neurons bordering the valve.

Having established our method to provide local mechanical stimuli, we characterized *Hydra’s* sensitivity to local touch and the associated neural response. We performed experiments using transgenic *Hydra* expressing GCaMP6s in neurons^30^. When we delivered mechanical stimuli, we found bright calcium signals generated by a small number of neurons in the hypostome and body column and a striking co-activation of many neurons in the ectodermal peduncle nerve ring (Figure 2b, Video 2, 3). This nerve ring activity appeared as either a single bright calcium spike (or “contraction pulse”) or a volley of bright calcium spikes (or “contraction burst”). We also found that the calcium-sensitive fluorescence averaged over a region of interest (ROI) surrounding the peduncle faithfully represented the contraction pulses and bursts measured from individual neurons (Figure 2 - Figure Supplement 2). When we analyzed single neuron calcium dynamics from this peduncle nerve ring, we found extremely high correlated activity as previously reported for contractions pulses and bursts (Figure 2 - Figure Supplement 2)^30, 31^. Given the similarity of these data between large and small ROIs, we chose to use the peduncle ROI to measure neuronal contraction bursts and pulses because it does not require single-neuron tracking, which significantly increased the throughput of our data analysis. We further confirmed that this signal is not the result of motion artifacts by measuring fluorescence from *Hydra* (nGreen) that express GFP pan-neuronally using a similar ROI. In that case we did not see the strong fluorescence signals associated with contraction pulses and bursts (Figure 2 – Figure Supplement 3e-f). Body length proved to be an unreliable quantification of contractions due to the stimulation artifacts (Figure 2 - Figure Supplement 3d); however, we were able to accurately measure muscle activity associated with contractions by imaging calcium spikes in the epithelial muscle cells (Figure 2 - Figure Supplement 4, 5 and Video 4, 5). Based on these experiments, we define *Hydra’s* “mechanosensory response” as calcium spikes in neural activity from the peduncle ROI and the associated calcium spikes in the epithelial muscles from the whole body if they occur within one second of mechanical stimulation onset (Figure 2c, d and Figure 2 – Figure Supplement 6).

Using the neuronal fluorescence calcium imaging described above, we found that the probability of the mechanosensory response depends on the intensity of the stimulus, which is consistent with many psychometric functions (Figure 2e). *Hydra* were five times more likely to contract within one second of receiving a strong mechanical stimulus than during a random one second interval without a stimulus (stimulus valve pressure 20 and 25 psi; response probability = 0.60 ± 0.06 and 0.77 ± 0.08, mean ± SEM, respectively; no stimulus valve pressure = 0 psi; response probability = 0.11 ± 0.01, mean ± SEM; Figure 2e, h). During mild stimuli, there was a slight increase (∼ 2x) in response probability above the spontaneous activity, although this increase was not statistically significant compared to spontaneous contraction bursts or pulses (valve pressure 10 and 15 psi; response probability = 0.23 ± 0.01 and 0.24 ± 0.03, mean ± SEM, respectively; Figure 2e, h). We found that *Hydra* did not respond to a weak mechanical stimulus that corresponded to a valve pressure of 5 psi (response probability = 0.09 ± 0.02, mean ± SEM; Figure 2e, h).

Further analysis of the calcium activity pattern revealed that the single contraction pulses (calcium spikes in the peduncle neurons) were more frequent when we repeated mechanical stimulation every 31 seconds for one hour. While spontaneous contraction pulses or bursts were observed roughly once every minute in microfluidics, when we stimulated *Hydra* with a strong mechanical stimuli, we found that the frequency nearly matched the 31 sec between stimuli (Interval between spontaneous contraction bursts or pulses = 72.98 sec ± 4.58, mean ± SEM; 66.68 ± 5.73 sec median ± SE; Interval between stimulated contraction bursts or pulses = 20 psi, 38.28 ± 1.77 sec, mean ± SEM; 31.14 ± 2.21 sec, median ± SE; 25 psi, 32.20 ± 1.04 sec, mean ± SEM; 31.14 ± 1.30 sec, median ± SE; Figure 2f,h). While the majority of spontaneous calcium spikes formed bursts, stimulated calcium spikes were roughly three times more likely to be a single contraction pulses (Percentage of spontaneous spiking activity that is a single contraction pulse = 0psi, 16.87 ± 5.38 % mean ± SEM; Percentage of stimulated spiking activity that is a single contraction pulse = 20 psi, 56.84 ± 2.48%; 25psi, 62.50 ± 5.23 %, mean ± SEM; Figure 2g, h).

### *Hydra* sensitivity to mechanical stimuli is lowest near the aboral end

Because of the diffuse neural architecture of *Hydra*, we expected each patch of *Hydra* tissue to be equally responsive to mechanical stimuli. However, when we stimulated transgenic *Hydra* expressing GCaMP6s (N = 3 whole animals, stimulated 40 times per region with 22 psi) at three different regions along their body (oral, middle body and aboral, stimulated 40 times at each region with 22psi), we found the aboral end of *Hydra* to be less sensitive than the center of the body (aboral region response probability = 0.1 ± 0.025; mid-body region response probability = 0.42 ± 0.025; mean ± SEM; p < 0.01; Figure 2 - Supplement figure 7a). Epitheliomuscular calcium imaging (N = 8 whole animals expressing GCaMP7b in ectodermal epitheliomuscular cells, stimulated 40 times in body column region with 22psi) also showed that the sensitivity to mechanical stimulation generally decreases towards the aboral end of the *Hydra* (Figure 2 – Supplement figure 7b). Furthermore, we found that the difference in sensitivity along the body was not an artifact due to differently sized *Hydra* experiencing different pressures from the microfluidic valves. We observed no statistically significant trend between animal size and response probability (Figure 2 – Supplement figure 7c). These findings, combined with the transcriptional analysis and in-situ hybridizations that indicate higher density of sensory neurons in the oral half of the *Hydra*^48^, suggest the oral end may be more sensitive to mechanical stimuli. Unlike other organisms that have unique motor responses like reversals or acceleration depending on the location of mechanical stimuli (e.g. *C. elegans*)^56, 57^, we observed that the same motor program was initiated regardless of where on the body the animal was touched. The only difference we observed was the fact that the response probability depended on where along the oral and aboral axis we delivered the mechanical stimulus.

### *Hydra*’s mechanosensory response is mediated by electrically-coupled cells

Based on the latency of the mechanosensory response, we hypothesized that sensorimotor information is transmitted by electrical activity in the *Hydra* and not by passive calcium diffusion through ectodermal cells. This hypothesis is supported by the fact that aneural *Hydra,* which no longer spontaneously contract in absence of stimuli, are capable of aversive contractile response to touch, however, their responses are slow and require strong mechanical stimulus^54, 55^. To further test our hypothesis, we created a transgenic *Hydra* strain that expresses the calcium indicator GCaMP7b (under the EF1ɑ promoter, see “Methods”) in the ectodermal epitheliomuscular cells. With this transgenic line we measured contraction pulses and contraction bursts by averaging calcium activity in all epitheliomuscular cells (Figure 2 – Figure Supplement 3, 5). We hypothesized that if the mechanosensory response was primarily mediated by calcium diffusion through epitheliomuscular cells, we would expect to see propagation of calcium activity from the site of the stimulation. However, this was not the case; we observed fast propagation of calcium activity throughout the entire ectoderm. Imaging calcium activity in the peduncle neurons showed that changes in neural activity correlated with the observed changes in epithelial muscle cells with increasing stimulus intensity (Figure 2 and Figure 2 – Figure Supplement 5). This suggests that the mechanosensory response is indeed mediated by the electrically-coupled cells. Although *Hydra*’s behavioral responses are slow compared to other invertebrates, the 0.5 sec average response time in our data could not be explained by calcium diffusion alone, which would take ∼100 sec to travel the average distance of ∼0.5-1 mm between the stimulation site and the peduncle or hypostome (Figure 2 – Figure Supplement 8a). Our data show the calcium signals from the peduncle neurons or ectodermal muscles start increasing within ∼0.1-0.2 sec following stimulation onset, reaching peak fluorescence at ∼0.5-0.6 sec regardless of the stimulus intensity (Figure 2 – Figure Supplement 8b-e). We found no difference in the calcium response times between neural and muscle calcium imaging, though this could be influenced by the dynamics of the calcium indicator, which typically cannot give information about latencies less than 50 ms^17^.

### Aboral neurons are not necessary for mechanosensory response in *Hydra*

Having quantitatively established *Hydra’s* sensorimotor response to mechanical stimuli, we next asked if specific regions of the nervous system play primary roles in mediating this response. Patch-clamp electrophysiology of individual neurons in *Hydra* has thus far been unsuccessful despite attempts by many research groups, and minimally invasive, cell-type specific neuromodulation techniques such as optogenetics have yet to be developed for *Hydra*. However, the animal’s regenerative abilities allow us to resect large portions of tissue without killing the animal, thus we can borrow from the tradition of lesioning brain regions to study their functions^52, 58–61^.

Because the *Hydra* body plan is radially symmetric with cell types primarily varying along the oral-aboral axis of the body column (Figure 1), we chose to make axial cuts across the body column to remove select neuronal populations from the animal. Our rationale was that these resections would remove entire or nearly entire groups of neuronal cell types. We then allowed ∼6-12 hours for the animal to recover. This recovery time helps to reduce the confounding contributions from initial tissue regeneration and allows animals to recover enough to tolerate microfluidic immobilization. This period is long enough to allow the wounds to close, the molecular response to injury to be completed^62, 63^, and the initial molecular events of regeneration to start; but it is *not* long enough for the animal to regenerate lost neurons, which takes approximately 30-72 hours (Figure 3 – Figure Supplement 1)^37, 61, 64, 65^. We confirmed that 6-12 hours after resection the animals indeed showed loss of specific neuronal cell types by measuring the expression levels of subtype-specific neuronal markers via qPCR (Figure 3 - Figure Supplement 2). To limit the stress from microfluidic immobilization that could exacerbate the resection wounds and affect activity, we shortened the duration of these experiments for the majority of the animals (40 min total, 20 min of no stimulation, 20 min of stimulation - valve on 1 sec at 22 psi or 0 psi, off 30 sec). Only three animals per each condition (stimulated and non-stimulated, and 5 different resections) were experimented on with a longer duration protocol as used previously (100 min total, 20 min no stimulation, 60 min of stimulation - valve on 1 sec at 22 psi or 0 psi, off 30 sec, 20 min no stimulation; Figure 3 - Figure Supplement 3).

We began resection studies by removing the peduncle and basal disk to create a “footless” *Hydra.* We hypothesized that aboral neurons may be important for coordinating and enhancing body contractions (Figure 3). We based this hypothesis on the fact that aboral neuron activity has a strong correlation with body contractions^30, 31^. In addition, the neuropeptide Hym-176C has been shown to induce ectodermal muscle contractions and is selectively expressed in the ectodermal peduncle neurons^48, 66–68^. Finally, the presence of gap junction protein innexin-2 in aboral neurons could facilitate fast electrical conductions that allows these neurons to fire synchronously^48, 55^. This could be necessary for enhancing neuromuscular signaling for body contractions. Because “footless” *Hydra* lacked peduncle neurons that we had used previously to measure contraction pulses and bursts (Figure 3 - Supplement Figure 3d-f, and Video 6-9), we performed these experiments using a transgenic *Hydra* line expressing GCaMP7b in the ectodermal epitheliomuscular cells, which allowed us to measure contraction pulses and bursts by averaging calcium activity in all the epitheliomuscular cells (using whole frame ROI which is more robust to motion artifacts than peduncle ROI; Figure 2 - Figure Supplement 3, Figure 3a,b, Figure 3 – Figure Supplement 3).

**Figure 3:**
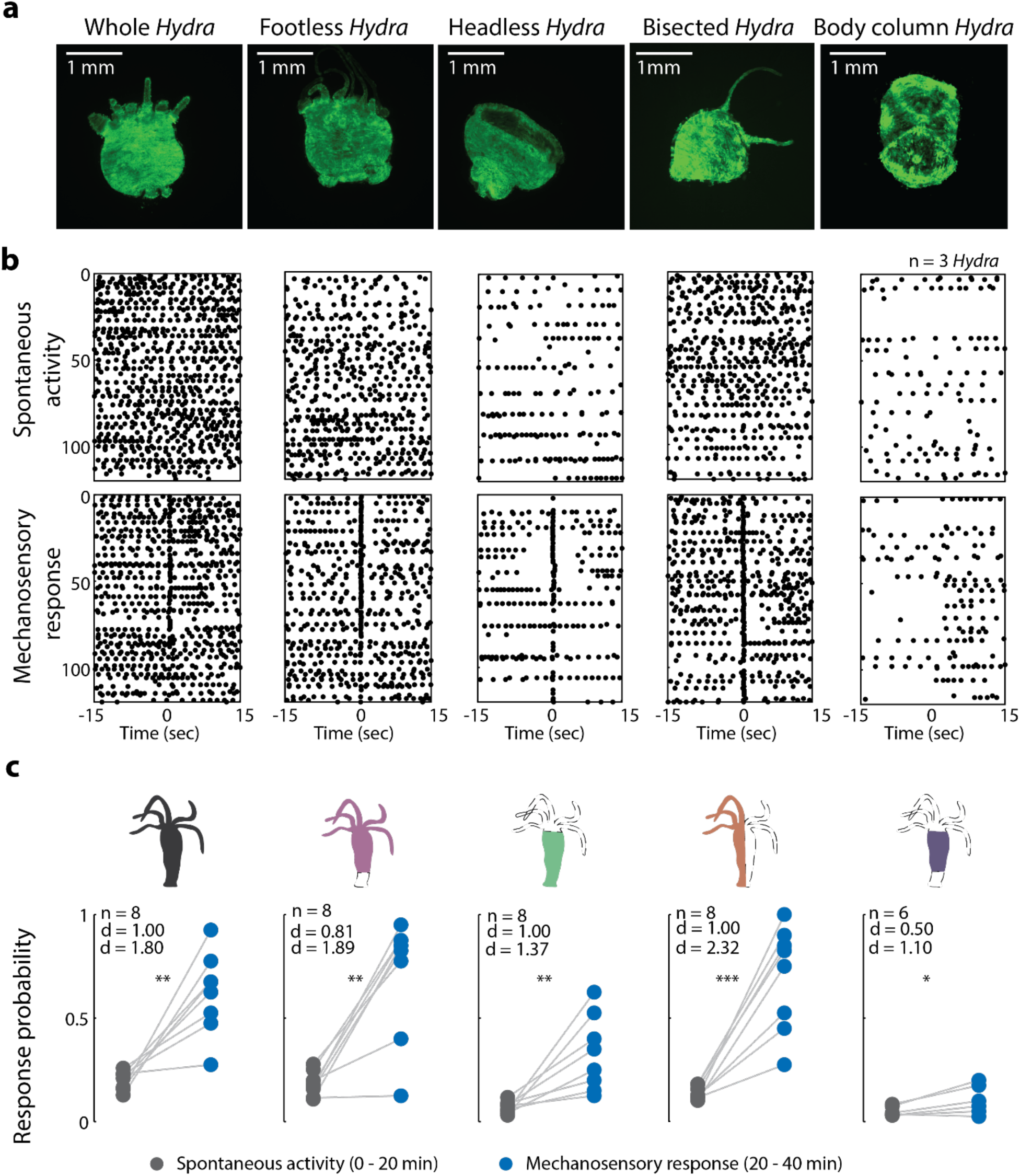
Oral region is important for mechanosensory response. a) Representative images of resection preparations (6-12 hours post-resections) of transgenic *Hydra* during contraction (GCaMP7b, ectodermal epitheliomuscular cells): whole (or control), footless, headless, bisected and body column animal. Entire frame ROI was used for analysis of the whole-body epithelial calcium activity. b) Representative raster plot of stimulus time aligned calcium spikes from three animals with multiple trials superimposed. Each black dot is a peak in calcium fluorescence identified as a contraction pulse. c) Response probability, fraction of trials that have at least one calcium spike (also contraction pulse) occurring within 1 sec of stimulation onset. Gray dots are the mean contraction probability during no stimulation (t = 0-20 min) calculated from 1 sec window shifted by ∼0.3 sec over 30 sec intervals. Blue dots are contraction probability during stimulation (t = 20-40 min) calculated from 1 sec response window during valve ‘on’. Light gray lines connect the probabilities for spontaneous contraction and mechanosensory response for each individual. Cartoon schematics of *Hydra* indicate the resections performed. Excised body regions are outlined with a dashed line and unfilled area. Color filled body regions indicate the portion of *Hydra* retained for the experiment. P-values from a paired t-test indicated as follows: n.s. = not significant; * = p < 0.05; ** = p < 0.01; *** = p < 0.001. Source data for the mechanosensory response in resected animals are available in the Figure 3 – source data 1.

Because neurons in the foot fire synchronously with body contractions, we expected “footless” animals to show significant changes in contraction behaviors (calcium spiking activity) to be significantly affected by their removal^30, 31, 69^, but this was not what we observed. Surprisingly, our experiments with “footless” animals showed that the aboral nerve ring was not required to regulate spontaneous contraction bursts or pulses or mechanosensory responses. After we removed the peduncle network in *Hydra*, we found the increase in contraction burst or pulse activity with stimuli (or mechanosensory response) in “footless” individuals was similar to the increase in activity observed in whole individuals (“Footless” N = 8 animals stimulated 20 min, Cohen’s d = 1.89, Cliff’s delta = 0.81, p < 0.01); Whole N = 8 animals stimulated 20 min, Cohen’s d = 1.80, Cliff’s delta = 1.00, p < 0.01; Fig 3b,c). Furthermore, we found no significant difference in either the spontaneous contraction probability or the mechanosensory response probability of “footless” animals and whole animals (“Footless” N = 3 animals not stimulated, spontaneous contraction probability = 0.13 ± 0.01, mean ± SEM; “Footless” N = 3 animals stimulated for 60 min, mechanosensory response probability = 0.88 ± 0.05, mean ± SEM) (Whole N = 3 animals not stimulated, spontaneous contraction probability = 0.15 ± 0.01, mean ± SEM; Whole N = 3 animals stimulated for 60 min, mechanosensory response probability = 0.73 ± 0.07, mean ± SEM; Figure 3 - Figure Supplement 3b,c, Video 4, 5,10, and 11).

### Oral neurons play a major role in mechanosensory response in *Hydra*

While the mechanosensory response in *Hydra* remained unaffected with the removal of the aboral nerve ring, we found that removal of the hypostome and tentacles (or “headless” *Hydra*) resulted in significant changes in both the mechanosensory response and spontaneous contraction bursts or pulses. When we measured the mechanosensory response in “headless” *Hydra,* we found that the animals still responded to mechanical stimulation with a significant increase in their contraction bursts or pulses, however, they did so with a lower magnitude compared to whole and “footless” individuals (“Headless” N = 8 animals stimulated for 20 min, p < 0.01, Cohen’s d = 1.37, Cliff’s delta = 1.00; Figure 3b, c). Specifically, the “headless” *Hydra* responded with (> 2x) lower probability compared to whole animals, and they also showed a (> 3x) lower probability of spontaneous contraction bursts and pulses (“Headless” N = 3 animals stimulated for 60 min, mechanosensory response probability = 0.29 ± 0.02, mean ± SEM; “Headless” N = 3 animals not stimulated, spontaneous contraction probability = 0.05 ± 0.01, mean ± SEM; Figure 3 - Figure Supplement 3b, c, Video 12 and 13).

To verify the reduction in mechanical response probability was specific to removing the oral network of neurons, and not simply the result of injury, we performed experiments with animals that we cut longitudinally along the body axis (“bisected” *Hydra*) to remove approximately the same amount of tissue while preserving the neuronal subtypes in both the oral and aboral networks. We found that these longitudinally “bisected” animals showed mechanosensory responses that were not significantly different from that of the whole, “headless”, or “footless” animals (“Bisected” N = 3 animals stimulated for 60 min, mechanosensory response probability = 0.54 ± 0.14, mean ± SEM; Figure 3 - Figure Supplement 3b, c). However, the magnitude of increase in contraction bursts or pulses activity due to stimulation in “bisected” individuals, though lower than whole and “footless” individuals, was larger than “headless” individuals (“Bisected” N = 8 animals stimulated for 20 min, p < 0.001, Cohen’s d = 2.32, Cliff’s delta = 1.0). This suggests that our observations in “headless” *Hydra* indeed depended upon the types of neurons removed during the headless resection and not simply an injury response. We also found that these longitudinally “bisected” animals had a lower probability of spontaneous contraction bursts or pulses than the whole animals but higher than “headless” animals. This suggests that the entire network needs to be intact for normal contraction bursts or pulses activity, and the loss of roughly half the network leads to some reduction in contractile activity (“Bisected” N = 3 animals not stimulated, spontaneous contraction probability = 0.08 ± 0.02, mean ± SEM; Figure 3 - Figure Supplement 3b, c Video 14 and 15).

We next asked if the body column alone is sufficient to mediate the mechanosensory response. To answer this question, we completely removed both oral and aboral regions. In “body column” animals, we found significant reduction in both the mechanosensory response and spontaneous contraction bursts and pulses relative to whole animals. The “body column” animals had a mechanosensory response probability that was not different from the mechanosensory response in “headless” animals, while the spontaneous contraction bursts and pulses probability was lower compared to that of “headless” animals (“Body column” N = 3 animals stimulated for 60 min, mechanosensory response probability = 0.19 ± 0.08; “Body column” N = 3 animals not stimulated, spontaneous contraction probability = 0.02 ± 0.01 mean ± SEM; Figure 3 - Figure Supplement 3b,c). Although we found a significant increase in contraction bursts and pulses with stimulation as compared to spontaneous contraction bursts and pulses activity in “body column” individuals similar to all resections, the magnitude of the increase was the lowest in “body column” animals (even lower than “headless” animals) (“Body column” N = 6 animals stimulated for 20 min, p < 0.05, Cohen’s d = 1.10, Cliff’s delta = 0.50; Figure 3b, c, Video 16 and 17). Moreover, we did not observe significant increases in contraction bursts and pulses over the same time period (comparing activity from 0 - 20 min with activity from 20 - 40 min, see “Methods”) in non-stimulated animals, suggesting the higher probability of contraction bursts and pulses was in fact due to mechanical stimulation (Figure 3 – Figure Supplement 4). Thus, based on the comparison between the probability of spontaneous contraction bursts and pulses and mechanosensory response in all resections, we found that the “body column” animals had a weak response to touch despite their slightly increased contraction bursts and pulses probability with mechanical stimulation.

### Oral and aboral networks show different patterns of activity during spontaneous contractions compared to mechanically stimulated contractions

To identify how the activity of neurons in the oral and aboral networks coordinate spontaneous and stimulated responses, we manually tracked the calcium activity of several neurons in the oral and aboral regions (20 minutes no stimulation, 10 minutes stimulation 1 sec every 31 sec, n = 3 *Hydra*; (Figure 4a, b, Figure 4 – Figure Supplement 1 and 2). We found there were at least two independent networks of neurons based on a correlation analysis (Figure 4c). Specifically, we time-aligned the calcium activity with either spontaneous contractions or mechanical stimulation events to classify these groups of neurons based on their activity (Figure. 4d, e, g). One group of correlated neurons found throughout the entire body showed averaged calcium activity less than one second after a mechanical stimulus and spontaneous activity that is consistent with previously reported Contraction Burst (CB) neurons^30^. We plot calcium dynamics of these CB neurons as shades of blue in Figure. 4 (and Figure 4 - Figure Supplement 1 and 2). These neurons show bursts of activity that are synchronized with muscle contractions and show calcium activity that is highly correlated with the average peduncle ROI. In addition to these CB neurons, we found other groups of correlated neurons with average calcium activity that is independent of the CB network. One group showed a distinctive pattern of activity following mechanical stimulation, but no distinctive activity associated with spontaneous contractions. Specifically, this group of neurons near the oral end responded approximately 10 seconds after mechanical stimulation (Figure. 4e, g). We found these putative “mechanically responsive (MR) neurons” in all three *Hydra* we analyzed and plot their calcium dynamics as shades of red in Figure. 4 (and Figure 4 - Figure Supplement 1 and 2). The fact that these MR neurons do not show activity associated with spontaneous contractions clearly indicates that they are not a part of the CB network, but rather these two distinct networks (CB and MR) are involved in the *Hydra’s* response to mechanical stimulation. We also note that these MR neurons do not fire periodically as would be expected for the Rhythmic Potential (RP) network^30^. In addition to CB and MR neurons, in *Hydra* 2 (Figure 4 - Figure Supplement 1) and 3 (Figure 4 - Figure Supplement 2) we also found neurons that were not associated with either the MR network or the CB network. These neurons we labeled as “unspecified groups” do not appear to be a part of the CB or RP networks previously characterized nor the MR neurons we identify here.

**Figure 4:**
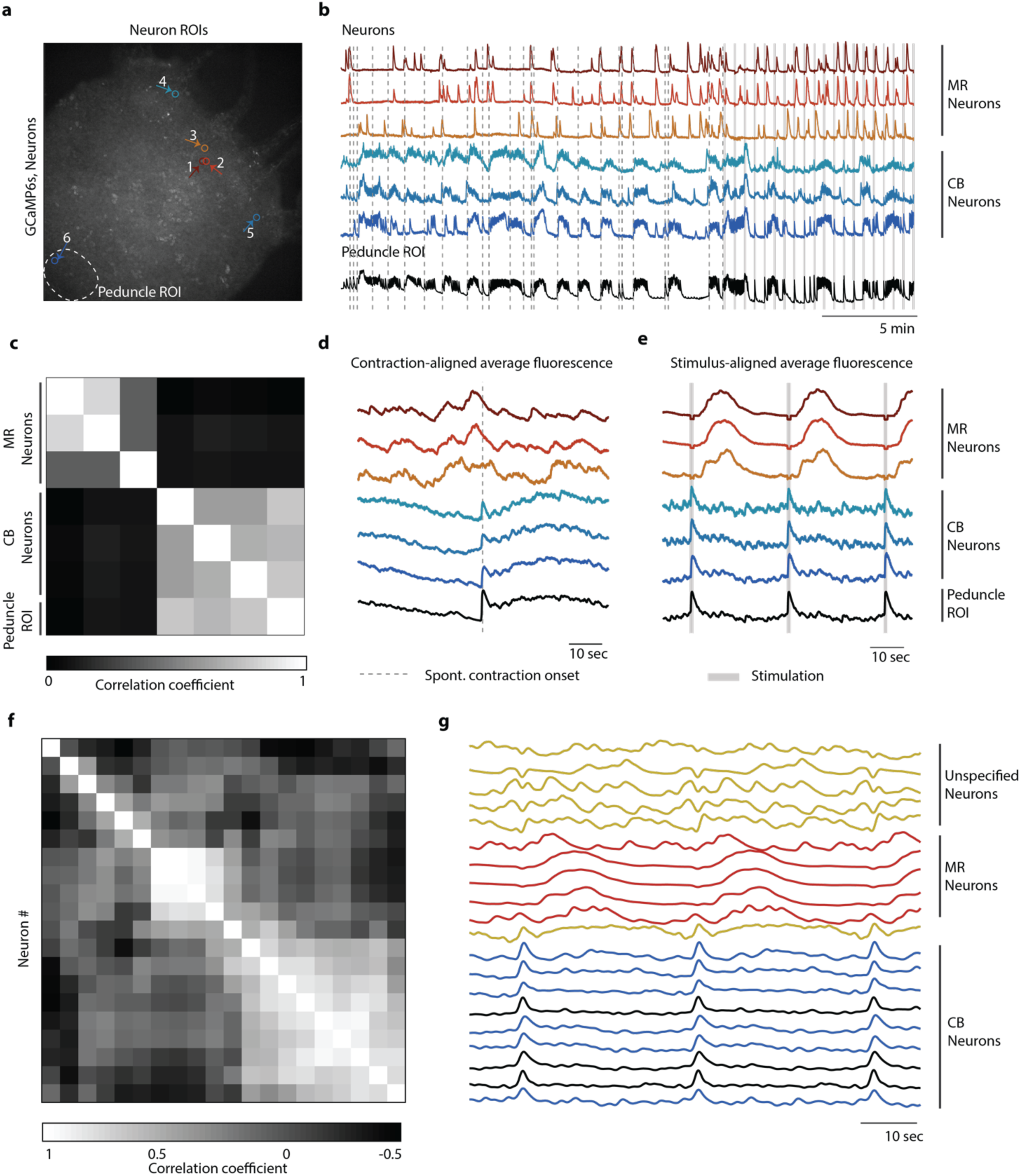
Distinct networks of neurons involved in spontaneous and stimulated behaviors. a) Fluorescent image of transgenic *Hydra* expressing GCaMP6s pan-neuronally. Individually tracked neurons ROIs are indicated by arrows, and peduncle ROI is outlined with a white dashed circle. b) Calcium fluorescence traces from single neurons (top 6 traces) and average calcium fluorescence from peduncle ROI (bottom trace). Mechanically responsive (MR) neurons are shown in shades of red. Contraction burst (CB) neurons are shown in shades of blue. c) Heat map shows the correlation coefficients of individually tracked neurons and peduncle ROI from *Hydra* 1, with color bar at the bottom. Correlation was computed using the entire 30 min calcium fluorescence from single neurons shown in (b). d) Average calcium fluorescence traces from each of the neurons and peduncle ROI during spontaneous behaviors time-aligned with the onset of spontaneous body contractions. Dashed line indicates the onset of body contraction. e) Average calcium fluorescence traces from each of the neurons and peduncle ROI during stimulated behaviors time-aligned with the onset of mechanical stimulation. Gray shaded rectangle indicates mechanical stimulation. f) Heat map shows the correlation coefficients of g) the average calcium fluorescence traces from each of the neurons and peduncle ROI (traces in black) from three different *Hydra* during stimulated behaviors time-aligned with the onset of mechanical stimulation. Three groups of neurons were identified across the three *Hydra.* CB neurons (traces in blue), MR neurons (traces in red) and unspecified neurons (traces in yellow). a-e) Calcium fluorescence traces and correlation analysis (entire 30 min of activity, 20 min of spontaneous activity and 10 min of stimulated activity) from one representative *Hydra*. f-g) Correlation analysis of the average stimulus-aligned calcium fluorescence (30 sec stimulation interval average from 10 min of stimulated activity) of each neuron pooled multiple *Hydra* (N = 3 *Hydra*) to highlight the three groups of neurons identified: CB, MR and unspecified. Source data for fluorescent calcium activity from single neurons are available in the Figure 4 – source data 1.

These data suggest that there are at least two separate pathways involved in the mechanosensory response. The first involves the CB neurons and muscle contractions. The second network involving the MR neurons responds more slowly, with a latency of several seconds. Because the activity of the MR neurons occurs after the contraction, their role in the behavioral response remains unclear.

## Discussion

Our experiments with “footless”, “headless”, “bisected”, and “body column” animals show that mechanosensory and spontaneous behaviors are regulated by neural ensembles that are localized to select regions of the animal; however, some properties of the mechanosensory response may be evenly distributed throughout the body. We demonstrate that localized touch produces an increased calcium activity in both peduncle neurons (Figure 2) and ectodermal epitheliomuscular cells (Figure 3), which is associated with body contractions (Figure 2 - Figure Supplement 4)^31^. We identified at least two neuronal networks (MR and CB network) with distinct neuronal activities mediating the stimulated responses (Figure 4), where the CB network of neurons show fast calcium responses, and the MR neurons show slower calcium response. It is possible that the MR neurons, primarily found in the oral end (N = 3 *Hydra*), may consist of ec4 neurons, which are the only neuronal subtype in the oral end with an unknown function. The other orally located ectodermal neuron populations, ec1B and ec3C, are suggested to belong to the CB and RP1 circuits, respectively^48^. *Hydra’s* responsiveness (i.e., probability of response to a mechanical stimuli) depends on the stimulus intensity (Figure 2). Interestingly, we found that the “headless” and “footless” animals can still respond to mechanical stimuli despite missing an entire regional neuronal network; however, “headless” animals show reduced responsiveness (Figure 3 and Figure 3 - Figure Supplement 3). The “body column” animals missing both regional neuronal networks have the most dramatically reduced responsiveness, suggesting that these two neuronal networks work together and play compensatory roles in mediating the mechanosensory response.

Surprisingly, although the activity of the peduncle nerve ring is strongly associated with spontaneous contractions, these neurons are not necessary for body contraction and response to mechanical stimulation. This raises the question of what role the peduncle nerve ring plays in *Hydra* behavior. One possible explanation is that this nerve ring coordinates body contractions by enhancing the neural signal to ectodermal cells. An additional 4-second reduction in body contraction duration in “body column” animals compared to headless animals supports the idea that the peduncle nerve ring is also involved in coordination of contractile behavior (Figure 3 – Figure Supplement 5). Furthermore, calcium activity propagates from the foot to the hypostome in whole animals during body contractions^70^, supporting a hypothesis that the peduncle network of neurons may be motor neurons.

Based on these observations, we propose a simple model for sensorimotor information flow in *Hydra* where we consider the hypostome as an integration point where sensory and motor information converge. The information is then communicated to the peduncle where it is amplified for coordinated whole body control. There may be sufficient functional redundancy between the hypostome and peduncle regions such that removal or damage to one of them is well tolerated in *Hydra*. Moreover, the diffuse network in the body column may retain minimal processing capabilities needed for weak sensorimotor responses. The fast and slow calcium responses from the CB and MR neurons, respectively, indeed support the hypothesis that *Hydra* have multiple, separate pathways for behavioral response.

Overall, the quantitative characterization of *Hydra’s* sensorimotor responses reported here help to build the foundation for a more comprehensive investigation of information processing in *Hydra*—an animal with clear advantages to supplement commonly studied model organisms in neuroscience. Important next steps include developing mechanistic models to describe sensory information processing that supports our results. Although our experiments describe general information flow in *Hydra*, having a cellular level control of neuronal activity would be a clear advantage for revealing the function of neuronal cell types as well as their functional connectivity in the sensorimotor circuits. We expect additional future work with optogenetic manipulation of specific neuronal subtypes combined with fast, volumetric and ratiometric imaging techniques will provide a more comprehensive approach for characterizing this sensorimotor processing in *Hydra*. Recognizing that calcium fluorescence imaging is limited in its ability measure single spikes^71^, we also expect other activity sensors (such as voltage indicators) will reveal what role the MR neurons play in behavioral response. Building upon the work reported here, one can then interrogate the roles of these regional networks with multiple sensory modalities, such as light and heat, to answer questions about how diffuse nervous systems may be capable of centralized information processing and multisensory integration. These studies may help reveal a comprehensive model for how internal states and external stimuli shape the behavioral repertoire in an organism with a highly dynamic neural architecture.

## Materials and Methods

### *Hydra* strains and maintenance

*Hydra* were raised in *Hydra* medium at 18°C in a light-cycled (12 hr: 12 hr; light: dark) incubator and fed with an excess of freshly hatched *Artemia nauplii* (Brine Shrimp Direct, Ogden, UT, #BSEP8Z) three times a week (protocol adapted from Steele lab). All experiments were performed at room temperature with animals starved for 2 days.

The transgenic line nGreen, kindly provided by Rob Steele, was generated by microinjecting embryos with a plasmid containing the *Hydra* actin promoter driving GFP expression^48^. The transgenic strains expressing GCaMP6s under the actin promoter in neurons and in ectodermal epitheliomuscles (Addgene plasmid: #102558) were developed by microinjections of the embryos by Christophe Dupre in the Yuste lab (Columbia University)^57^. The transgenic strain expressing GCaMP7b in ectodermal epitheliomuscular cells was co-developed by Juliano Lab (University of California, Davis) and Robinson Lab (Rice University). Briefly, the plasmid with codon optimized GCaMP7b under the EF1a promoter was constructed by GenScript (www.GenScript.com). Injections were performed as previously described^72^ with the following modifications: 1) injection solution was prepared by mixing 1 µL 0.5% phenol red (Sigma P0290-100ML) with 6 µL plasmid DNA solution prior to centrifugation, and 2) embryos were fertilized for 1-2 hours prior to injection. Plasmid promoters were cloned in expression vector pHyVec2 (Addgene plasmid: # 34790) using restriction sites Nsil. Plasmids were prepared by Maxiprep (Qiagen, Valencia, CA) and eluted in RNase-free water. A plasmid DNA solution of 1.4 µg/uL was injected into embryos using an Eppendorf FemtoJet 4x and Eppendorf InjectMan NI 2 microinjector (Eppendorf; Hamburg, Germany) under a Leica M165 C scope (Leica Microscopes, Inc; Buffalo Grove, Il). Viable hatchlings with mosaic expression were propagated by asexual reproduction, and asexual buds were screened and selected for increasing amounts of transgenic tissue until a line was established with uniform expression in the ectodermal epithelial cells.

### Fluorescence imaging of *Hydra* nerve net

Distribution of neurons in the *Hydra* nerve net was fluorescently imaged with transgenic *Hydra vulgaris* expressing GFP (nGreen) in neurons and neuronal progenitors (Figure 1a)^48^. *Hydra* was anesthetized with 0.05% chloretone and immobilized in a ∼160 µm tall microfluidic chamber^31^. High-resolution fluorescence imaging was performed using a confocal microscope (Nikon TI Eclipse) and 10x (0.45NA) objective, where the *Hydra* was imaged at a single plane with multiple fields of views stitched together to obtain an image of the whole animal (Figure 1a).

### *Hydra* resections

*Hydra* were placed in a petri dish and covered with enough *Hydra* medium to prevent desiccation. When *Hydra* were relaxed and stationary, resections were performed with a single incision with a scalpel across the body. We referenced published images of in-situ hybridizations of different cell types and spatial expression patterns to guide the location of incisions. For “footless” *Hydra,* an axial cut above the peduncle removed ∼one third of the lower body, which included the peduncle and the basal disk. For “headless” *Hydra,* an axial cut below the hypostome removed ∼one third of the upper body, including the tentacles and the hypostome. For “bisected” *Hydra*, a transverse cut starting from the tip of the hypostome to the basal disk was made along the midline of the body. For “body column” *Hydra*, an axial cut above the peduncle removed the lower body followed by another axial cut below the hypostome to remove the upper body region. This preparation resulted in an open tube body. *Hydra* were stored in an 18°C incubator after the excisions and until beginning the experiments. *Hydra* can seal open wounds within ∼3-4 hours and repopulate the neuronal population to regain functionality in 30-72 hours (Figure 3 – Figure Supplement 1, see “Imaging regeneration of peduncle network”). To allow *Hydra* time to recover but not regain the functionality of lost neuronal cell types, we performed experiments 6-12 hours post amputation.

### Imaging regeneration of peduncle network

Transgenic *Hydra* (GcaMP6s, neurons) were axially cut in the middle of the body column to generate an oral and aboral halves (Figure 3 – Figure Supplement 1a). The oral half was immobilized between two coverslips with a ∼110 µm PDMS spacer. Calcium fluorescence was conducted for 20 min every 2 hr on Nikon TI Eclipse inverted microscope with 20% excitation light from Sola engine and GFP filter cube. We captured frames at ∼10Hz (100ms exposures) with Andor Zyla 4.2 sCMOS camera with NIS software. We used 4x (0.2 N.A) objective for wide-field imaging to fit the entire *Hydra* in the field of view reduce the likelihood of *Hydra* migrating out of the imaging frame. *Hydra* were exposed to blue excitation light for 20 min during imaging and remained under dark conditions for 100 min between subsequent imaging timepoints. One animal was imaged for ∼20 hours starting with 1 hr post resection (Figure 3 – Figure Supplement 1b). *Hydra* had a visibly open wound at the first imaging point but was not detectable after 3 hr. There were no visibly active neurons in the regenerating aboral end, indicating that the resected peduncle neurons had not regenerated. Another animal was imaged for ∼30 hours starting with 37 hr post resection (Figure 3 – Figure Supplement 1c). At the first imaging time point (t = 37 hr post resection), there were few neurons in the peduncle region that were active during body contractions, and these groups of neurons resembled a nerve ring as early as 41 hr post resection. Although the peduncle nerve ring seemed to have formed at this point, it appeared to not be as densely populated qualitatively as observed in whole animals.

### Microfluidic device fabrication

The mechanical stimulation devices are double-layer microfluidic devices with push down valves custom-designed with CAD software (L-edit) and fabricated using standard photo- and soft-lithography techniques. All master molds were fabricated with transparency photomasks and SU-8 2075 (MicroChem). The master mold for the bottom *Hydra* layer (circular chambers, 3 mm diameter) was fabricated with the height of ∼ 105-110 µm thick pattern (photoresist spun at 300rpm for 20 sec, 2100 rpm for 30 sec). The master mold for the top valve layer (9 individual circular valves, 3 x 3 arrangement, 400 µm diameter each) was fabricated with height of ∼110 µm thick (photoresist spun at 300rpm for 20sec, 2100 rpm for 30 sec). Polydimethylsiloxane (PDMS) Sylgard 184 was used to cast microfluidic devices from the master molds. The bottom *Hydra* layer (10:1 PDMS spun at 300rpm for 40 sec ∼3 hr post mixing; cured for 12 min at 60°C) was bonded to the valve layer (∼4 mm thick, 10:1 PDMS; cured for ∼40 min at 60°C with holes punched for inlet ports) with oxygen plasma treatment (Harrick Plasma, 330 mTorr for 30 sec) and baked for at least 10 min at 60°C. *Hydra* Insertion ports were hole-punched through the two layers for *Hydra* layer with 1.5 mm biopsy punches, and the devices were permanently bonded (O2 plasma treatment, 330 mTorr for 30 sec) to 500 µm thick fused silica wafer (University Wafers) and baked for at least 1 hr at 60°C. The design files for the photomasks and step-by-step fabrication protocols will be available on www.openHydra.org (under Resource hub/Microfluidics).

*Hydra* were immobilized and removed from the microfluidic device using syringes to apply alternating positive and negative pressures as previously reported^31^. The microfluidic devices were reused after cleaning similarly to the protocol previously reported. Briefly, the devices were flushed with deionized water, sonicated (at least 10 min), boiled in deionized water (160°C for 1 hr), and oven dried overnight.

After repeated use, the PDMS stiffness can change and affect the valve deflection and the actual force experienced by *Hydra* through the PDMS membrane. Additionally, uncontrollable conditions during the device fabrication process can also lead to small differences between devices. As a result, all data for Figure 2 was taken with a single device and the response curve was used to calibrate (identify pressure that yielded ∼ 60% response probability equivalent to 20-22 psi stimulus intensity) new devices.

### Microfluidic mechanical stimulation

We used compressed air to inflate the microfluidic valves. For temporal control over valve (on/off) dynamics, we used a USB-based controller for 24 solenoid pneumatic valves (Rafael Gómez-Sjöberg, Microfluidics Lab, Lawrence Berkeley National Laboratory, Berkeley, CA 94720) and a custom-built Matlab GUI (available at www.openHydra.org) that allowed setting the stimulation parameters, such as the duration of valve ‘on’ (1 sec), duration of valve ‘off’ (30 sec), and the duration of stimulation (60 min, 119 cycles of valve ‘on’ and ‘off’), and pre, post-stimulation acclimation/control period (20 min). We used a pressure regulator to manually control the air pressure into the valve manifold. In summary, we set the stimulation pressure with a pressure gauge to regulate the flow of air into the valve manifold. This valve manifold was controlled with a USB valve controller that allowed us to programmatically inflate the valve (turn it ‘on’) with pressurized air with custom stimulation parameters.

To test sensory motor response to mechanical stimuli, we used pressurized air to inflate the push-down valve and cause it to press down on the *Hydra* immobilized in the bottom layer. Each experimental condition had at least three *Hydra* each. For a given condition, replication experiments were conducted on different days. After an animal was immobilized inside the *Hydra* chamber, we selected one valve (from the nine valves over the entire chamber) that was directly above the mid-body column region to deliver stimuli. Although *Hydra* were free to move, we did not observe large displacement most of the time and, as a result, the same valve remained in contact with the animal throughout the stimulation period.

We adjusted the air pressure using a pressure regulator for each experiment, and the valves were inflated using a USB valve controller (see above). The full-length stimulation experiment consisted of 20 min of no stimulation (control/acclimation) followed by 60 min of stimulation period (except habituation experiment where the stimulation period was 120 min), where valves were pulsed with constant pressure (0 (control), 5, 10, 15, 20, 22 or 25 psi) for 1 sec every 31 sec, then another 20 min of no stimulation (control/acclimation). Shorter stimulation experiment (used for whole-animal muscle imaging of various resections) consisted of 20 min of no stimulation (acclimation/control) followed by 20 min of stimulation period (valves pulsed for 1 sec every 31 sec with a constant pressure of 0 or 22psi). We chose a 20 min initial control period based on the high sensitivity to abrupt changes in light intensities (especially to blue wavelengths used for excitation of GCaMP) in *Hydra* which leads to increased contractile activity for 2-5 min. Even with the stimulus repeated for two hours at a constant inter-stimulus interval, we found no obvious evidence of sensitization, habituation, or stimulus entrainment in *Hydra* (Figure 2 – Figure Supplement 9). As a result, we chose not to randomize the inter-stimulus interval.

### Distribution of mechanical forces

We characterized the distribution of force exerted by a microfluidic valve by quantifying the movements of neurons due to mechanical stimulation. We performed fluorescence imaging in transgenic *Hydra* (nGreen) expressing GFP in neurons for ∼8 min. We captured 8,000 frames at ∼16Hz (50 msec exposures) with 4x objective (0.2 N.A.) and Andor Zyla 4.2 sCMOS camera with 2×2 binning (1024×1024 frame size) using MicroManager. *Hydra* was stimulated in the middle of the body column (valve pulsed for 1 sec every 31 sec, 5 times).

To quantify the movement of neurons and tissue throughout the *Hydra* body, we tracked a total of 222 neurons that were visible for 1 min capturing the first two stimulation trials. We performed semi-automated tracking with TrackMate plugin (ImageJ/(Fiji)^72–74^ and manually corrected the tracks where neurons were misidentified. From these tracks, we calculated the displacement of each of the neurons between each frame (∼50 msec). The average cellular displacement (0.4 µm per frame, 50 msec) was calculated by averaging the cellular displacements from all frames when the valve was not pressurized. We found a significantly increased displacement (6.2 um, p < 0.001) just after the valve was pressurized (and after the valve was depressurized 1 sec later). We then generated a vector map of neuronal displacements by calculating the change in position of each of the neurons in the frame just after the valve was pressurized for the first stimulation trial (Figure 2 - Figure Supplement 1a). We also plotted the cellular displacement for each of the neurons and the location of those neurons relative to the center of the valve. We averaged the highest three displacements in 50 µm radial band increments from the valve center to quantify how far the mechanical forces extended. This was a more conservative measurement as the neurons in different tissue layers (endodermal and ectodermal layers furthest from the valve) may have experienced different forces. By taking the average of the highest three displacements in 50 µm radial bands increments from the valve center, we identified the majority of the (shear) force was experienced by neurons bordering the valve in 250 µm radius.

### Fluorescence intensity from GFP *Hydra*

To prove that the fluorescence intensity changes we observed with calcium imaging are due to calcium activity and not motion artifacts from body contractions, we compared the average fluorescence intensities from three different transgenic *Hydra* lines: 1) expressing GFP pan-neuronally (nGreen line), 2) expressing GCaMP6s pan-neuronally, and 3) expressing GCaMP7b in ectodermal epitheliomuscular cells (Figure 2 - Figure Supplement 3). For each *Hydra*, we compared the average fluorescence intensities from three different regions of interest (ROIs) that included the peduncle region, whole frame (for whole body) region, and valve region. We also measured the body length by taking the major axis of an ellipse fitted along the oral-aboral axis of the body column (from apex of the hypostome to the basal disc) after binarizing the fluorescence image.

For this analysis, we used fluorescence imaging from transgenic *Hydra* expressing GFP pan-neuronally (nGreen) performed in “Distribution of mechanical forces”. *Hydra* was stimulated in the middle of the body column (valve pulsed for 1 sec every 31 sec, 5 times). We used calcium imaging from transgenic *Hydra* (expressing either GCaMP6s pan-neuronally or GCaMP7b in ectodermal epitheliomuscular cells) stimulated in the middle of the body column (valve pulsed for 1 sec every 31 sec, 120 times). The average fluorescence intensity and body length traces were time-aligned to the onset of mechanical stimulation to show 1) changes in average fluorescence intensity in peduncle ROI and the whole frame ROI are due to calcium activity (increase in fluorescence from in GCaMP lines) and not motion artifact (no change in fluorescence from GFP line) and 2) increase or decrease in body length or average fluorescence from valve ROI are affected by stimulation artifacts.

### Average calcium fluorescence from large and small ROIs in the peduncle

We performed fluorescence imaging in transgenic *Hydra* expressing GCaMP6s in neurons for ∼1 min during spontaneous behaviors (*Hydra* was not stimulated). We captured frames at ∼16Hz (50 msec exposures) with a 10x objective (0.45 N.A.) and Andor Zyla 4.2 sCMOS camera with 3×3 binning (682×682 frame size) using MicroManager.

Because calcium fluorescence decreases when neurons are inactive, tracking multiple neurons in a highly deformable region is a challenge. Nonetheless, we tracked 9 neurons from the peduncle nerve ring that were visible (enough to track) for 1 min. We performed semi-automated tracking with TrackMate plugin (ImageJ/(Fiji)^72–74^ and manually corrected the tracks where neurons were misidentified. We also used a large peduncle ROI, similar to how we used an ROI for quantifying neural response to stimulation in all other experiments. We calculated the average calcium fluorescence trace from small ROIs for individual neurons and large ROI for the peduncle region. Fluorescence intensity was normalized by calculating ΔF/F, where ΔF = F-F_0_ and F_0_ is the minimum fluorescence from prior time points. The large peduncle ROI had highly correlated calcium fluorescence activity with smaller ROIs for individual neurons which are known to belong to the contraction burst circuit (Figure 2 - Figure Supplement 2). As a result, for all other experiments we use large peduncle ROI when measuring the neuronal activity.

### Whole-animal imaging of neural activity at different stimulus intensities

We characterized the neural response to repeated local mechanical stimuli at 0 (control), 5, 10, 15, 20 and 25 psi pressure with animals that expressed GCaMP6s in neurons (Figure 2). Each pressure condition was experimented with three animals on different days using the same device and stimulation protocol (valve pulsed for 1 sec every 31 sec for 60 min) to generate the pressure-response curves (total of 18 animals). Calcium fluorescence imaging for all experiments was conducted for the 100 min duration of the stimulation protocol (see Mechanical Stimulation subsection) on Nikon TI Eclipse inverted microscope with 20% excitation light from Sola engine and GFP filter cube. We captured 100,000 frames at ∼16Hz (50 msec exposures) with Andor Zyla 4.2 sCMOS camera with 4×4 binning using MicroManager. For imaging neural activity, 12-bit low noise camera dynamic range was used. To synchronize the stimulation onset times with imaging, we use used a data acquisition device (LabJack U3) to record the TTL frame out signal (fire-any, pin #2) from the Zyla and the valve on/off timestamps from the valve controller.

We used 4x (0.2 N.A.) objective for wide-field imaging to fit the entire *Hydra* in the field of view to reduce the likelihood of *Hydra* migrating out of the imaging frame. There are neurons in the oral region (hypostome) that are co-active with aboral region (peduncle) neurons during body contractions; however, neurons in the hypostome appeared to be much sparser than those in peduncle (Video 18). Discerning these hypostomal network neurons required higher magnification which significantly reduced the field of view such that considerable amount of nervous tissue could move in and out of the imaging plane (z-plane) or frame (xy-plane), making it difficult to obtain reliable calcium fluorescence time-series.

### Whole-animal imaging of neural activity with different stimulation regions

To map the sensitivity of different body parts to mechanical stimuli, we performed calcium imaging of multiple animals (N = 3, whole animals expressing GCaMP6s in neurons) using imaging settings previously described with modified experimental protocol. Based on the range animal sizes (1-2.5 mm) and the size of the valve (400 um), we stimulated *Hydra* body at three different regions along the oral-aboral axis: 1) the oral end (upper third of the body), 2) the aboral end (lower third of the body), and 3) the third near the mid-body column. Each animal was stimulated at three different locations (20 min no stimulation, stimulation region #1 - 1 sec every 31 sec for 20 min, ∼2 min no stimulation, stimulation region #2 - 1 sec every 31 sec for 20 min, ∼2 min no stimulation, stimulation region #3 - 1 sec every 31 sec for 20 min using 22 psi). We randomized the order in which the three different body regions were stimulated in each of the three animals to avoid any stimulus entrainment artifacts. We then analyzed the peduncle nerve ring activity in response to mechanical stimulation of different body regions as details in “Analysis of calcium activity” and “Analysis of mechanosensory response”.

### Whole-animal imaging of neural activity in resected animals

For experiments that involved body lesions (Figure 3 – Figure Supplement 3), the same experiment protocol and imaging settings as previously described (see “whole animal imaging of neural activity at different stimulus intensities”, 20 min no stimulation, stimulation 1 sec every 31 sec for 60 min, 20 min no stimulation) were used with either 0 psi (control) or 20 - 22 psi (∼ 60% response probability). Each condition group had three animals (total of 18 animals). The “headless” and “bisected” animals were prepared with the appropriate body regions removed as described previously (*‘‘Hydra* resections”). Due to difficulty in tracking neurons and weak GCaMP expression in body column neurons, we were unable to quantify responses from “footless” and “body column” *Hydra* and thus excluded them from experiments.

### Simultaneous electrophysiology and calcium imaging of ectodermal epitheliomuscular cells

Electrical activity from the epitheliomuscular cells was measured simultaneously with calcium imaging of the ectodermal epitheliomuscular cells in transgenic *Hydra* using a nano-SPEARs device previously reported (Figure 2 – Figure Supplement 4)^31^. Briefly, transgenic *Hydra* expressing GCaMP6s in the ectodermal epitheliomuscular cells starved for at least 48 hr were used to measure the activity of the muscles (10 fps, 30 min, 4x objective with 15% light intensity). An inverted microscope with GFP filter and Andor Zyla 4.2 were used for capturing images. All electrical data was obtained with an Intan Technologies RHD2132 unipolar input amplifier (http://intantech.com) at a sampling rate of 1KHz, low frequency cutoff and DSP filter of 0.1 Hz and high frequency cutoff of 7.5 KHz. From the calcium activity traces, we identified 30 sec intervals of either high amplitude activity or low amplitude activity to perform cross-correlation analysis. The high and low amplitude activity regions were manually identified with a threshold of 20% of the highest peak in the calcium activity. The high amplitude activity region occurred during contraction bursts. The low amplitude activity region occurred during tentacle contractions for muscular activity imaging. Both the Intan amplifier and the Zyla were triggered with the same TTL signal. However, to account for any offset in the timing of the electrical and optical data, we measured the maximum of the cross-correlogram in a 50 ms (approximately one duty cycle of the trigger signal) window rather than the cross-correlation at zero offset to generate the correlation maps. For correlation map, each frame was down sampled to 64 x 64 pixels, and the fluorescence trace for the downsampled pixels across the 30 sec intervals was cross-correlated with electrical activity during the same 30 sec interval. The intensity of color in the correlation map was used to indicate correlation values.

### Whole-animal imaging of epitheliomuscular activity at different stimulus intensities

We characterized the muscle response to mechanical stimuli using animals that expressed GCaMP7b under the EF1a promoter in ectodermal epitheliomuscular cells (Figure 3). We measured contraction pulses and contraction bursts by averaging calcium activity in all the epitheliomuscular cells (Figure 3a). The correlation between peduncle nerve ring activity and muscle contractions has been previously established based on simultaneous electrophysiology and neuronal^31^ or ectodermal epitheliomuscular cells calcium imaging (Figure 2 – Figure Supplement 4). The imaging protocol for epitheliomuscular cells was similar to the imaging of neural activity (see “whole animal imaging of neural activity at different stimulus intensities”), except a 16-bit camera dynamic range was used to avoid saturating the sensor.

We first developed a partial psychometric curve for calcium activity of epitheliomuscular cells to confirm the dependence of response on stimulus intensity. Three animals were imaged for 20 min without stimulation and then stimulated at three different pressures (20 min at 15psi, 20 min at 20 psi, 20 min at 25psi; or in reverse order), and the epitheliomuscular response curve was used to identify the stimulus intensity (∼ 22psi) which yielded ∼ 60% response probability (Figure 2 - Figure Supplement 5). This stimulation intensity was also selected for stimulation of N = 8 whole animals (40 min of imaging; 20 min of no stimulation, 20 of stim for 1 sec every 31 sec with 22 psi) and animals with different body regions removed.

### Whole-animal imaging of epitheliomuscular activity in resected animals

The various body regions were removed as described previously (*‘‘Hydra* resections”) to prepare “headless”, “footless”, “bisected” and “body column” *Hydra*. Note, this transgenic line of *Hydra* expressing GCaMP7b in the ectodermal cells showed some signs of deficit. They were particularly sensitive to being handled and were more likely to dissociate after ∼30 min in the chambers during long-term microfluidic immobilization and fluorescence imaging. The resected *Hydra* were especially difficult to image for the entire 100 min without any cell dissociation. This could be due to the specific promoter used for driving GCaMP expression or that phototoxicity is higher when there is a high expression of GCaMP in a larger number of cells. Because the lesioned animals were more likely to be damaged during microfluidic immobilization, the mechanical stimulation protocol (see Mechanical Stimulation subsection) was shortened to total of 40 min of imaging with 20 min of no stimulation followed by 20 min of repeated stimulation (1 sec every 31 sec for 20 min). For three animals per each resection (total of 30 animals), we used the full-length protocol (100 min of imaging: 20 min of no stimulation, 60 min of stimulation every 31 sec, and 20 min of no stimulation) to obtain higher statistical power for quantifying response probability (Figure 3 – Figure supplement 3). A total of 6-8 animals were stimulated for each resection condition (whole, N = 6 not-stimulated, N = 8 stimulated; “footless”, N = 4, not-stimulated, N = 8 stimulated; “headless”, N = 5 not-stimulated, N = 8 stimulated; “bisected”, N = 5 not-stimulated, N = 8 stimulated; “body column” N = 3 stimulated, N = 6 stimulated; total 61 animals, Figure 3; 30 of which were experimented with long (100 min) stimulation protocol Figure 3 – Figure supplement 3).

### Analysis of calcium activity

To analyze neural response, we used the average fluorescence from the peduncle region. Tracking individual neural responses was difficult due to high deformability of the body and lack of fluorescence markers when neurons were not active; as a result, we looked at the synchronous firing activity of the neurons in the peduncle region, which is known to have high correlation with body contractions. To analyze epitheliomuscular response, we used average fluorescence from the whole animal to obtain the fluorescence signal over time and analyzed these signals similarly to neural response. Briefly, using ImageJ (Fiji)^72, 73^ we used a ROI over the peduncle region or the whole *Hydra* to obtain fluorescence signal over time for neuronal or epitheliomuscular calcium activity, respectively.

From the fluorescence signal, we detected the large calcium spikes as individual contractions (contraction pulses or bursts) using MATLAB peak finding algorithm. We generated a raster plot of calcium spiking activity time-aligned with the stimulus where each row represented one stimulation trial with 15 sec before and after stimulation onset. These raster plots were used to calculate the probability for spontaneous contraction and mechanosensory response. For contractile behavior analysis, we then annotated the calcium spikes. Single calcium spikes were labeled as single pulse contraction. A volley of calcium spikes was labeled as a contraction burst. Both of these led to behavioral contractions, thus time between contractions was calculated as the time between the end of a contraction event (single pulse or burst) and the start of the next one (as shown in Figure 3 – Figure Supplement 5). Percent of contractions that are single pulse contractions was calculated as the fraction behavioral contractions that were single calcium spikes (not bursts). Contraction duration was used to indicate the amount of time contraction behavior lasted (time from rise in fluorescence signal to return to baseline, as shown in Figure 3 – Figure Supplement 5)

### Analysis of mechanosensory response

To obtain the response probability (whether an animal had a contraction pulse—either a single pulse or a pulse from a burst) at the time of stimulation (when valve was pressurized), we generated a raster plot of fluorescence spikes time-aligned to stimulus onset and superimposed for each 30 sec intervals (time between stimulus). We used an hour of activity (t = 20-80 min) to generate the raster plot and calculate response probability. We defined the 1 sec window while the valve was pressed as the response window for each trial (Figure 2 – Figure Supplement 6). We then calculated the fraction of all trials (119 trials over 60 min or 40 trials over 20 min) per animal that had at least one fluorescence spike (“contraction pulse”) in the 1 sec response window following stimulus onset to obtain the contraction probability. Extraction of raw fluorescence for ROIs was performed with ImageJ (Fiji)^72, 73^ and post-processing analysis was performed with MATLAB (using peak-finding algorithm to detect spikes). A one-way ANOVA with Bonferroni correction was used for statistical analysis.

For animals with shorter experiment duration, the mechanosensory response probability in stimulated animals was calculated over the 20 min segment (time = 20-40 min) with a fraction of all trials (40 trials over 20 min) that had at least one fluorescence spike. A one-way ANOVA with Bonferroni correction was used for statistical analysis when comparing multiple conditions. A paired t-test was used to compare the difference between probability of spontaneous contraction and mechanosensory probability in the same animals for each of the conditions. The effect size (magnitude of difference) was measured with Cohen’s d and Cliff’s delta.

### Analysis of spontaneous contraction

For non-stimulated animals, fluorescence activity from one hour of activity (time = 20-80 min) was converted into a raster plot with multiple (119 trials over 60 min) 30 sec long intervals (to match the stimulation interval in stimulated animals). We obtained “stimulation times” using a DAQ to record the signal from valve controller except the air pressure was set to 0 psi. The spontaneous contraction probability was calculated by taking the average of response probability from a random 1 sec interval (Figure 3 and Figure 3 – Figure Supplement 3). Briefly, by sliding a 1 sec window by ∼0.3 sec over the x axis (30 sec of stimulation interval) on the raster plot (pooled from all three animals for Figure 4a, per individual stimulated animal for Figure 3 – Figure Supplement 3), we generated the distribution of the fraction of trials (119 trials x 3 animals over 60 min) with at least one contraction event in a random 1 sec response window (Figure 3 – Figure Supplement 3). The distributions were compared with a Kruskal-Wallis test.

For animals with shorter experiment duration, spontaneous contraction probability in stimulated animals was calculated similar to non-stimulated animals above, except the raster plot was generated with the first 20 min of fluorescence activity (time = 0-20 min) when no stimulation was applied. Briefly, we generated a distribution of random probabilities by sliding a 1 sec window over the x-axis on the raster plot and used the distribution mean as the spontaneous contraction probability to compare with mechanosensory response probability from the same animal with a paired t-test (Figure 3). For non-stimulated animals with shorter experiment duration, spontaneous contraction probability was also calculated over the second 20 min interval (t = 20-40 min) to confirm the increase in contraction probability during mechanical stimulation was in fact due to stimuli and not just from temporal variation in spontaneous contractions (Figure 3 – Figure Supplement 4). Note, these experiments were performed with *Hydra* constrained to ∼110um thick microfluidic chambers. Although animals are able to behave under such confinements, the range of behavioral motifs and rates may be altered due to compression.

### Neuron subtype gene expression analysis with RT-qPCR

Resected *Hydra* were prepared as described above, with 12 polyps per biological replicate (except 6 whole polyps for control) and a total of two biological replicates per treatment. Approximately 12 hrs post resections, tissue was frozen in 1 mL Trizol at -80°C until use. RNA was isolated using the Zymogen RNA Clean and Concentrator kit (Zymogen #R1017) with an in-column Zymogen DNAse I digestion (Zymogen #E1010) following the manufacturer’s protocol. cDNA was synthesized using 1 µg of purified RNA and Promega M-MLV RNase H Minus Point Mutant Reverse Transcriptase (Promega, Madison, WI M3682) using the manufacturer’s protocol for oligo dT-primed synthesis. cDNA synthesis was validated via PCR, and all cDNA samples were diluted 1:10 in nuclease-free water for use in qPCR experiments.

Each sample was run in three technical replicates per gene in a 10 uL qPCR reaction using Bio-Rad SsoAdvanced universal SYBR green master mix (Bio-Rad, Hercules, CA 1725271). Samples were run on a CFX96 Touch Real-Time PCR Detection System (Bio-Rad 1855195). Data were analyzed with the 2^^-ΔΔCt^ method using the ‘tidyverse’ package in R^75, 76^.

Briefly, Cq values from technical replicates were pooled for subsequent analyses. *rp49* was used as an internal control to calculate ΔCq values after first being found to give similar results across all treatments when compared to a second control gene, *actin.* All results were normalized to the “whole” animal samples. A template-free water control was performed for all primer sets to ensure contamination-free reactions. All qPCR primers (Table 1) were validated via a 10-fold serial dilution standard curve to have a binding efficiency over 90%.

**Table 1:**
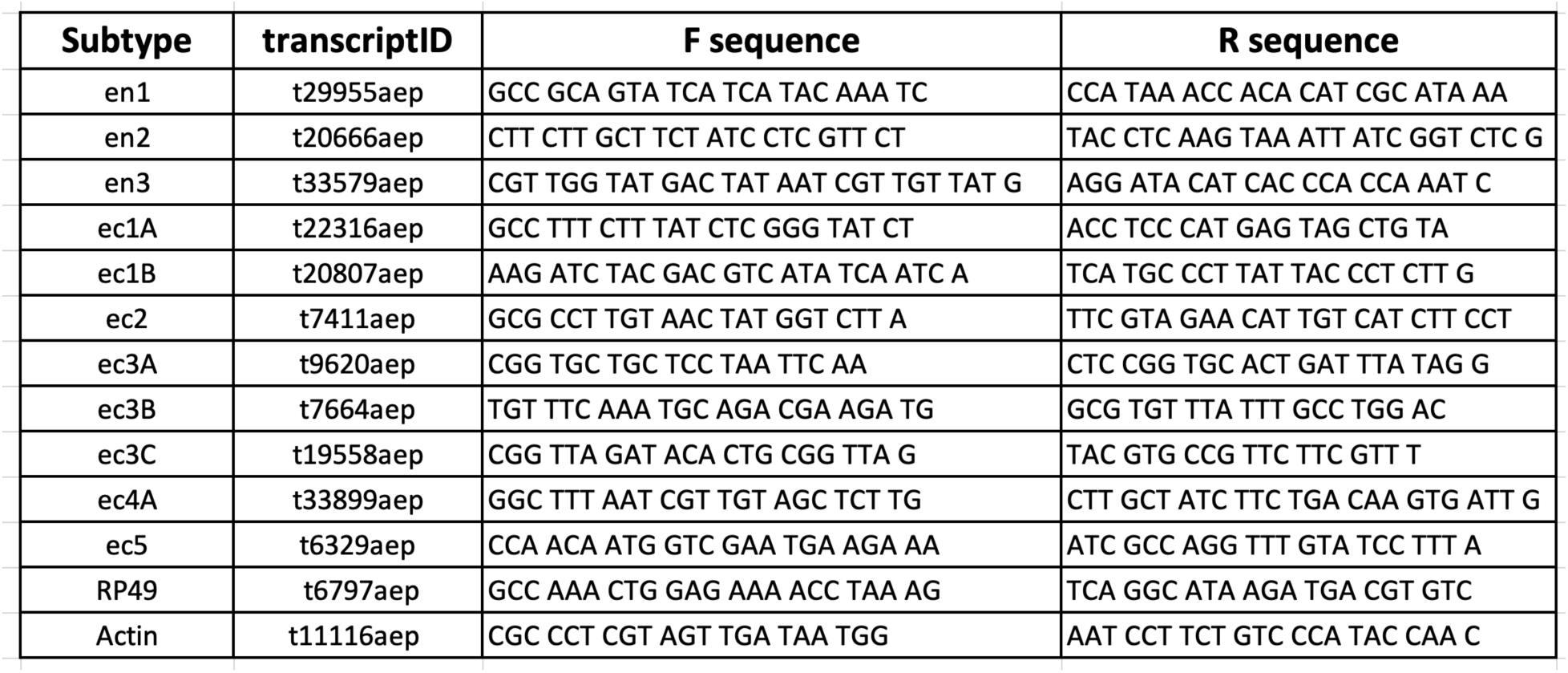
qPCR primers for neuronal subtypes.

### Whole animal imaging with single neuron resolution

We performed fluorescence imaging in transgenic *Hydra (N = 3)* expressing GCaMP6s in neurons for ∼30 min (30 min of imaging: 20 min of no stimulation, 10 min of stimulation every 31 sec). We captured frames at ∼16Hz (50 msec exposures) with a 10x objective (0.45 N.A.) and Andor Zyla 4.2 sCMOS camera with 3×3 binning (682×682 frame size) using MicroManager.

Because calcium fluorescence intensity is low (almost indiscernible from the autofluorescence of *Hydra*) when neurons are inactive, tracking multiple neurons in a highly deformable region without static nuclear fluorescence (such as RFP) is a challenge. Nonetheless, we manually tracked 5-6 neurons throughout the animal body that were visible (enough baseline fluorescence to track even when the neurons were inactive) for the entire duration of the imaging. We performed semi-automated tracking with TrackMate plugin (ImageJ/(Fiji)^72–74^ to track individual neurons with small circular ROIs (∼5 µm radius). We manually tracked (interpolating the current ROI location based on the past and future locations) the neurons when the fluorescence levels were dimmer than the background autofluorescence or when neurons were misidentified by TrackMate. We also used a large peduncle ROI, similar to how we used an ROI for quantifying neural response to stimulation in all other experiments. We calculated the average calcium fluorescence traces from small ROIs for individual neurons and large ROI for the peduncle region. Each of the fluorescence traces were corrected by calculating ΔF/F_0_, where F_0_ was the mean fluorescence intensity of the trace.

We performed the correlation analysis (MATLAB) of the calcium fluorescence time-series for the individual neuron ROIs and peduncle ROI to identify groups of neurons with correlated activity. We calculated the correlation coefficients for the individual neurons and the peduncle ROI by correlating the entire 30 min calcium fluorescence trace for each of the neurons. These correlation coefficients were shown to be statistically significant then randomly reshuffled calcium fluorescence. We divided each of the neuronal calcium time-series into 30 blocks. These individual blocks were randomly recombined to generate reshuffled time-series for each of the neurons to calculate correlation coefficients. This random reshuffling of each neuron was repeated 1000 times to obtain the mean correlation coefficients for each of the neuron pairs (Figure 4 – Figure Supplement 3d-f). We then calculated the z-score (x-μ)/σ, where x is the correlation coefficients of the original raw time-series, μ is the mean correlation coefficients of the time-series randomly reshuffled 1000 times and σ is the standard deviation of the correlation coefficients of the time-series randomly reshuffled (Figure 4 – Figure Supplement 3g-i). To identify the neuronal clusters, we used a hierarchical clustering algorithm (MATLAB function ‘linkage’ with Euclidian distance of correlation coefficients) on the correlation coefficients. We then used leaf order from the linkage tree to sort the neurons and generate the correlation heatmap.

We calculated the average fluorescence traces from the time-aligned calcium fluorescence to either spontaneous contractions or stimulus events to identify the roles of these neurons. For spontaneous contractions, the calcium fluorescence (t = 0-20 min) was time-aligned to the onset of contraction bursts or pulses from the peduncle ROI. For stimulated contractions, the calcium fluorescence (t = 20-30 min) was time-aligned to the onset of stimulus recorded with a DAQ (see “Whole-animal imaging of neural activity at different stimulus intensities”). We further confirmed the different neuronal clusters identified by performing correlation analysis on all neurons from all three Hydra by calculating the correlation coefficients for the average fluorescence traces from the time-aligned calcium fluorescence to stimulus events (Figure 4f.g).

## Acknowledgements

We thank Dr. Rob Steele (UC Irvine), Dr. Christophe Dupre (Harvard University), and Dr. Rafael Yuste (Columbia University) for sharing transgenic *Hydra*. We also thank Dr. Guillaume Duret and Dr. Caleb Kemere for useful discussions. The promoter sequence for EF1a for generating transgenic *Hydra* was provided by Jack Cazet (UC Davis, Juliano lab), and the plasmid with GCAMP7b was cloned (and codon optimized) by GenScript. The open source USB valve controller was built based on the design by Rafael Gómez-Sjöberg, Microfluidics Lab, Lawrence Berkeley National Laboratory, Berkeley, CA 94720. This work is funded by NSF EDGE. K.N.B. is funded by a training fellowship from the Keck Center of the Gulf Coast Consortia on the NSF IGERT: Neuroengineering from Cells to Systems 1250104. We also thank the Rice Shared Equipment Authority and nanofabrication cleanroom facility where the devices were fabricated.

## Figure Supplements

**Figure 2 – Figure Supplement 1: Distribution of mechanical forces.** Fluorescent image (grayscale) of transgenic *Hydra* (nGreen) expressing GFP in neurons and neural progenitors. The Oral region is on the left. Aboral//peduncle region is on the right. Arrows (cyan) overlayed indicate the displacement of individual cells immediately before and after the valve is pressurized for stimulation (∼50 msec). The lengths of the arrows are enlarged to be visible with the scaled size of the largest displacement ∼31 µm as indicated by the legend (lower left). Semi-transparent (yellow) circle indicates the location of the valve (400 µm diameter). b) Displacement of neurons due to stimulation varies with the location of the neurons from the valve center. Each semi-transparent blue dot represents a single neuron (n= 222 neurons). Black dots (connected by black dashed line) represent the average displacement of the highest three displacements within a 50 µm radial band. Microfluidic valve has a radius of 200 µm indicated by the yellow shaded region. Majority of the cumulative displacement (>60 %) is localized to a 250 µm radius bordering the valve indicated by the blue dashed line at 450 µm. Neurons more than 750 µm from the microfluidic valve center have negligible displacement (<6% cumulative displacement) indicated by the green dashed line.

**Figure 2 – Figure Supplement 2: Average calcium fluorescence from large peduncle ROI correlated with calcium fluorescence from smaller ROIs for individual peduncle neurons**. a) Fluorescent image of transgenic Hydra expressing GCaMP6s pan-neuronally. Dashed while circle indicates the peduncle ROI. Small cyan circles indicate the individual neuron ROIs. Peduncle region is in the top right. The oral end of the Hydra (not shown in the image) is oriented towards the bottom left corner. b) Average calcium fluorescence trace (black) from the large peduncle ROI. Average calcium traces (gray) from individual neurons in the peduncle.

**Figure 2 – Figure Supplement 3: Spikes in calcium fluorescence are due to calcium activity not motion artifacts.** a) Fluorescence image of transgenic *Hydra* (nGreen) expressing GFP in neurons false colored with hot green colormap. Neurons appear in white. b) Fluorescence image of transgenic *Hydra* expressing GCaMP6s in neurons false colored with hot green colormap. c) Fluorescence image of transgenic *Hydra* expressing GCaMP7b in ectodermal epitheliomuscular cells false colored with hot green colormap. a-c) Body length (white arrow), valve ROI (top circles), peduncle ROI (larger bottom circles) and whole frame ROI (entire frame square) annotated on all three images for comparison. d) Average body length trace time-aligned to stimulus onset from GFP *Hydra* (left) and GCaMP *Hydra* (middle, right). Light gray shaded region indicates when the stimulation is on. e) Average fluorescence trace calculated from the annotated peduncle ROI from GFP *Hydra* (left) and GCaMP *Hydra* (middle, right). f) Average fluorescence trace calculated from the whole frame ROI from GFP *Hydra* (left) and GCaMP *Hydra* (middle, right). g) Average fluorescence trace calculated from the annotated valve ROI from GFP *Hydra* (left) and GCaMP *Hydra* (middle, right).

**Figure 2 – Figure Supplement 4: Simultaneous electrophysiology and calcium imaging of ectodermal epitheliomuscular cells.** (a) Photograph (left) of a microfluidic immobilization chamber filled with green dye. Black box highlights the recording region in the microfluidic chamber (110 µm tall). False colored scanning electron micrograph (middle) shows the recording region (blue, photoresist; light grey, Pt; dark grey, silica) on the nano-SPEAR chip (50 µm tall). Inset shows a zoom-in of the Pt electrode (light grey) suspended mid-way between the top and bottom of the photoresist sidewall (blue). Brightfield image (right) shows *Hydra* immobilized in the microfluidic chamber placed on top of the nano-SPEAR chip with combined 160 µm tall recording region (Reproduced from Figure 2A, Badhiwala et al, 2018^31^ by permission of The Royal Society of Chemistry. This panel is not covered by the CC-BY 4.0 license, and further reproduction of this panel would need permission from the copyright holder.) b) Simultaneous electrophysiology and calcium imaging in transgenic *Hydra* (GCaMP6s, ectodermal epitheliomuscular cells). Top trace shows mean fluorescence (ΔF/F) from ectodermal epitheliomuscular cells (whole frame ROI) in *Hydra* (GCaMP, ectodermal) (10Hz). Bottom trace shows simultaneously recorded electrical activity from the *Hydra* (1KHz). High and low activity periods identified based on peak amplitude. Inset: left box shows correlation during high activity period, contraction bursts. Traces show peaks in fluorescence (top trace) coinciding with peaks in electrophysiology (bottom trace). A representative fluorescence image shows high levels of fluorescence thus calcium activity in the entire body during contraction burst events. Correlation map spatially plotting the correlation coefficient shows the entire body has calcium activity correlated with electrophysiology. Inset: right box shows correlation during low activity period, tentacle pulses. Traces show peaks in fluorescence (top trace) coinciding with peaks in electrophysiology (bottom trace). A representative fluorescence image shows high levels of fluorescence in the tentacle region during tentacle pulses. Cross correlation map shows the tentacle region has calcium activity correlated with electrophysiology. Almost all spikes in electrophysiology coincide with spike in fluorescence from ectodermal epitheliomuscular cells (and behavioral contractions).

**Figure 2 – Figure Supplement 5: *Hydra*’s epitheliomuscular response is dependent on the mechanical stimulus intensity**. a) Gray trace (top) is the stimulation protocol. 20 min no stimulation, 20 min of repeated stimulation (1 sec ‘on’, 30 sec ‘off’) at 15 psi, 20 min of repeated stimulation (1 sec ‘on’, 30 sec ‘off’) at 20 psi, and 20 min of repeated stimulation (1 sec ‘on’, 30 sec ‘off’) at 25 psi. Stimulus ‘on’ times indicated by vertical lines. Entire frame ROI used for analysis of the whole-body epithelial calcium activity. Representative calcium fluorescence trace (blue) from ectodermal epitheliomuscles (GCaMP7b) from animal stimulated with three different stimulus intensities for 20 min each (15, 20 and 25 psi). Representative calcium fluorescence trace (teal) from ectodermal epitheliomuscles (GCaMP7b) from animal stimulated with three different stimulus intensities for 20 min each (25, 20 and 15 psi). The decrease in fluorescence amplitude observed for both increasing in and decreasing stimulus intensity is due to photobleaching of the calcium indicator. b) Mechanosensory response probability at different stimulus intensities (N = 3 animals). Response probability, fraction of trials that have at least one calcium spike (also contraction pulse) occurring within 1 sec of stimulation onset.

**Figure 2 – Figure Supplement 6: Mechanosensory response window.** a) Different response windows tested (1 sec in blue; 5 sec in orange; 10 sec in yellow; and 15 sec in purple) (Top). The lengths of the rectangles correspond to time (sec) in the raster plot below. Stimulus trace in gray. Representative raster plot of time-aligned contraction pulses from multiple trials superimposed from one animal stimulated at 20 psi every 31 sec for 60 min (Bottom). Each black dot is a spike in calcium fluorescence identified as a contraction pulse. Stimulus is applied from 0 - 1 sec as indicated by a step in stimulus trace. b) Response probability, fraction of trials that have at least one calcium spike (also contraction pulse) occurring during different response windows (1 sec in blue; 5 sec in orange; 10 sec in yellow; and 15 sec in purple within stimulation onset).

**Figure 2 – Figure Supplement 7: Mechanical sensitivity of different body regions in *Hydra***. a) Response probability of transgenic Hydra (N=3 animals expressing GCaMP6s in neurons) stimulated at three different body regions: Oral, Mid-body and Aboral. Annotated Hydra below the plot indicates the three stimulation regions used. b) Response probability of transgenic Hydra (n=8 animal expressing GCaMP7b in ectodermal epitheliomuscular cells) Response probability is calculated using average calcium fluorescence from neurons in the peduncle ROI in a) and using average calcium fluorescence from the ectodermal epitheliomuscular cells from the entire body in b) and c).

**Figure 2 – Figure Supplement 8: *Hydra’s* mechanosensory response time is faster than passive calcium diffusion through epitheliomuscular cells** a) Distance calcium diffuses passively over given time (left, black line plot) approximated using passive diffusion equation (right). Light brown shaded region indicates the range of body length (∼ 0.5 - 1 mm) of *Hydra* in microfluidic chambers used for the experiments. Teal shaded region indicates the average time between stimulus onset and observed spike in fluorescence (mean response time 0.5 - 1 sec). Yellow shaded region is the theoretical mean calcium diffusion time calculated assuming passive diffusion. b-e) Mean calcium fluorescence time-aligned to stimulus and normalized. Stimulus is applied at 0 sec (valve ‘on’ from 0 - 1 sec). b) Mean calcium activity from the peduncle network of neurons at different stimulus intensities. c) Mean calcium activity from ectodermal epitheliomuscular cells at different stimulus intensities. b-c) The line colors correspond to different stimulus intensities shown at the bottom. d) Mean calcium activity from peduncle network of neurons in different body resections stimulated at 20 psi (pressure for ∼ 60% response probability in whole animals). e) Mean calcium activity from ectodermal epitheliomuscular cells in different body resections different body resections stimulated at 22 psi (pressure for ∼ 60% response probability in whole animals). d-e) The line colors correspond to the different resections shown at the bottom with cartoon schematic of the resected *Hydra* with the same color.

**Figure 2 – Figure Supplement 9: Long-term mechanical stimulation** a) Representative calcium fluorescence trace from the peduncle region from an animal (GcaMP6s, neurons) not stimulated and animal mechanically stimulated with 20 psi. Stimulus protocol in gray, 20 min no stimulation, 120 min of repeated stimulation (1 sec ‘on’, 30 sec ‘off’) at 20 psi followed by no stimulation. b) Spontaneous and stimulated (mechanosensory response) contraction pulse probabilities for the first and second hour of stimulation compared for each animal. Gray circles are mean contraction pulse probability during no stimulation calculated from an average of 1 sec window shifted by ∼ 0.3 sec over 30 sec intervals (N = 3). Blue circles are contraction probability during stimulation calculated from 1 sec response window during valve ‘on’ (N = 2). Light gray lines indicate the change in probabilities from hour 1 to hour 2 for each individual. (paired t-test, n.s. = not significant)

**Figure 3 – Figure Supplement 1: Regeneration of the peduncle network.** a) Summary schematic of the experiment performed. Dashed line indicates where the incision was made to remove the lower half of the transgenic *Hydra* body (GcaMP6s, neurons) to discover when the peduncle neuronal network is regenerated. b) Fluorescence images of “footless” *Hydra* during body contractions at various timepoints t = 1 hr, 5 hr and 9 hr after bisection. Pink arrow indicates the open wound visible at t = 1 hr but not at other timepoints. c) Fluorescence images of another “footless” *Hydra* during body contractions at various timepoints t = 37 hr, 39 hr, 41 hr. Blue dashed circle indicates the region where the peduncle network is found. Blue arrows indicate some of the neurons in the peduncle network with clearly well-connected neurons at t = 41 hr.

**Figure 3 - Figure Supplement 2: RT-qPCR analyses of neuron subtype-specific gene expression in resected animals demonstrates loss of specific neuron subtypes.** RT-qPCR was used to test for the loss of specific neuron subtypes in whole, tube (no head or foot), headless, footless, and bisected *Hydra* using uniquely expressed biomarkers for each subtype. There is not a specific biomarker for ec3A (located in the basal disk), so the marker used to test for the presence or absence of this cell type is also expressed in ec3B (located in the body column). Therefore, expression of the ec3A/B marker gene is reduced, but not completely lost in animals with resected feet (“tube” and “footless”). However, ec5 expression (located in the peduncle above the basal disk) is completely lost in animals with resected feet, thus it is clear that the ec3A subtype is completely lost in these animals. Biomarkers for neuron subtypes located in the head and tentacles (ec1B, ec2, ec3C, ec4) are completely lost in animals with resected heads (“headless” and “tube”). Data were analyzed with the 2^^-ΔΔCt^ method and results were normalized to the housekeeping gene *RP49* and to expression in whole animals.

**Figure 3 – Figure Supplement 3: Mechanosensory response from ectodermal epitheliomuscular cells and neurons in resected *Hydra*.** a) Representative images of resection preparations of transgenic *Hydra* (GCaMP7b, ectodermal epitheliomuscular cells: whole (or control), “footless”, “headless”, “bisected”, and “body column”. White dashed square indicates the whole frame ROI used for quantifying response. b) Spontaneous probability of at least one spiking event (or contraction pulse) occurring during a random 1 sec window. (1 sec window slide by ∼ 0.3 sec across 30 sec stimulation intervals; Kruskal-Wallis test with post hoc dunn-sidak correction) c) Mechanosensory response probability during 1 sec of valve on (Response probability, fraction of trials that have at least one calcium spike (also contraction pulse) occurring within 1 sec of stimulation onset). Large circles indicate average values from all animals combined for each condition. Small circles indicate probability from individual animals (one-way anova with post hoc Bonferroni correction). Error bars are standard error of mean (S.E.M.) (N = 3 *Hydra* for each condition; analysis from 60 min of stimulation which is imaging t = 20 – 80 min; n.s. = not significant, ** = p < 0.01, **** = p < 0.0001, ***** = p < 0.00001). d) Representative images of resection preparations of transgenic *Hydra* (GCaMP6s, neurons: whole (or control), “headless”, and “bisected”. White dashed circle indicates the peduncle ROI used for quantifying response. e) Spontaneous probability of at least one spiking event (or contraction pulse) occurring during a random 1 sec window. (1 sec window slide by ∼ 0.3 sec across 30 sec stimulation intervals; Kruskal-Wallis test with post hoc dunn-sidak correction) f) Mechanosensory response probability during 1 sec of valve on. Large circles indicate average values from all animals combined for each condition. (Response probability, fraction of trials that have at least one calcium spike (also contraction pulse) occurring within 1 sec of stimulation onset.) Small circles indicate probability from individual animals (one-way anova with post hoc Bonferroni correction). Error bars are standard error of mean (S.E.M.) (N = 3 *Hydra* for each condition; analysis from 60 min of stimulation which is imaging t = 20 – 80 min; n.s. = not significant, ** = p < 0.01, **** = p < 0.0001, ***** = p < 0.00001).

**Figure 3 – Figure Supplement 4: Contraction activity in non-stimulated animals.** Cartoon schematic of resection preparations of transgenic *Hydra* (GCaMP7b, ectodermal epitheliomuscular cells): whole (or control), “footless”, “headless”, “bisected” and “body column” animals. Entire frame ROI used for analysis of the whole-body epithelial calcium activity. Gray circles are mean contraction probability during t = 0 - 20 min. Blue circles are mean contraction probability during t = 20 - 40 min, (this time corresponds to when stimulated animals receive mechanical stimuli). Mean probability calculated from 1 sec window shifted by ∼ 0.3 sec over 30 sec intervals. (Response probability, fraction of trials that have at least one calcium spike (also contraction pulse) occurring within 1 sec of stimulation onset). Pressure in valves = 0 psi. Light gray lines pair the spontaneous contractions probability from the first 20 min interval with the second 20 min interval from each individual animal. (Whole N = 4, footless N = 3, headless N = 3, bisected N = 5, body column N = 3 *Hydra*, paired t-test, n.s. = not significant)

**Figure 3 – Figure Supplement 5: Hypostome and peduncle nerve rings work together to coordinate contractile behavior** a) Representative fluorescence trace used to calculate time interval between contractions. b) Time interval between spontaneous contractions in animals with different resections: whole, “footless”, “headless”, “bisected” and “body column”. c) Time interval between stimulated contractions in animals with different resections. d) Representative fluorescence trace used to calculate time interval between contraction pulses. For illustration purpose, only a select few of the time intervals are shown. e) Time interval between spontaneous contraction pulses in animals with different resections. f) Time interval between stimulated contraction pulses in animals with different resections. g) Representative fluorescence trace used to calculate contraction duration. h) Duration of spontaneous contractions in animals with different resections. i) Duration of stimulated contractions in animals with different resections. N= 3 *Hydra* (GcaMP7b, Ectodermal epitheliomuscles) per resection. (Kruskal-Wallis test with dunn-sidak correction, n.s. = not significant, ** = p < 0.01, *** = p < 0.001, **** = p < 0.0001,***** = p < 0.00001,****** = p < 0.000001)

**Figure 4 – Figure Supplement 1: Single cell correlation analysis.** a) Fluorescence image of transgenic *Hydra* (#2) expressing GCaMP6s pan-neuronally. Individually tracked neuronal ROIs are indicated with circles. The neurons are numbered in the order their traces are appear in c-e with the same colors. Peduncle ROI outlined with white dashed line. b) Calcium fluorescence traces from single neurons (top 6 traces) and average calcium fluorescence from peduncle ROI (bottom trace). Mechanically responsive (MR) neurons are shown in shades of red. Contraction burst (CB) neurons are shown in shades of blue. Unspecified neurons are shown in yellow, and their activity do not resemble any of the previously identified neuronal networks (contraction burst, rhythmic potential or the mechanically responsive reported here). c) Heat map shows the correlation coefficients of individually tracked neurons and peduncle ROI. Colorbar is at the bottom. d) Average calcium fluorescence traces from each of the neurons and peduncle ROI during spontaneous behaviors time-aligned with the onset of spontaneous body contractions. Dashed line indicates the onset of body contraction. e) Average calcium fluorescence traces from each of the neurons and peduncle ROI during stimulated behaviors time-aligned with the onset of mechanical stimulation. Gray shaded rectangle indicates mechanical stimulation.

**Figure 4 – Figure Supplement 2: Single cell correlation analysis.** a) Fluorescence image of transgenic *Hydra* (#3) expressing GCaMP6s pan-neuronally. Individually tracked neuronal ROIs are indicated with circles. The neurons are numbered in the order their traces are appear in c-e with the same colors. Peduncle ROI outlined with white dashed line. b) Calcium fluorescence traces from single neurons (top 5 traces) and average calcium fluorescence from peduncle ROI (bottom trace). Mechanically responsive (MR) neurons are shown in shades of red. Contraction burst (CB) neurons are shown in shades of blue. Unspecified neurons are shown in yellow, and their activity do not resemble any of the previously identified neuronal networks (contraction burst, rhythmic potential or the mechanically responsive reported here). c) Heat map shows the correlation coefficients of individually tracked neurons and peduncle ROI. Colorbar is at the bottom. d) Average calcium fluorescence traces from each of the neurons and peduncle ROI during spontaneous behaviors time-aligned with the onset of spontaneous body contractions. Dashed line indicates the onset of body contraction. e) Average calcium fluorescence traces from each of the neurons and peduncle ROI during stimulated behaviors time-aligned with the onset of mechanical stimulation. Gray shaded rectangle indicates mechanical stimulation.

**Figure 4 – Figure Supplement 3: Random shuffling of single cell correlation analysis.** Individual neuronal fluorescence time-series were randomly shuffled to show the correlation coefficients were not due to random chance in all three *Hydra* (each row is a different *Hydra*). Original correlation coefficients from 10 mins of raw fluorescence time-series (a-c), shuffled correlation coefficients from randomly reshuffled blocks reconstructing 10 mins of fluorescence time-series (d-e) and z-score of correlation coefficients calculated based on random reshuffling (g-i). Original correlation coefficients were calculated from the 10 mins of raw fluorescence activity for each of the neurons during mechanical stimulation. Shuffled correlation coefficients were calculated by taking the mean of correlation coefficients from randomly reshuffled fluorescence time-series. Briefly, the raw fluorescence time-series from 10 mins of mechanical stimulation was divided into 100 blocks, which were then randomly recombined for each of the neurons to calculated correlation coefficients and this random reshuffling was repeated 1000 times. Z-score was calculated using the mean and standard deviation of the correlation coefficients calculated from random reshuffling repeated 1000 times. The heatmap for z-score (g-i) resemble the original correlation coefficients (a-c) indicating the correlation coefficients of the raw fluorescence time-series were not a result of random chance. Except for the diagonals which represents autocorrelation, the mean correlation coefficients resulting from randomly reshuffled fluorescence (e-f) are zero for all neuron pairs indicating the correlation due to random chance.

## Description of Videos

**Video 1:**

Microfluidic system to study mechanosensory response in *Hydra*. Dashed blue circle indicates the microfluidic valve that presses down on *Hydra.* Mechanosensory response from neurons and epitheliomuscular cells is shown.

**Video 2:**

Spontaneous neural calcium activity in normal animals. Dashed blue circle indicates the ROI used for calcium trace shown in blue (bottom). (Playback 100x.)

**Video 3:**

Stimulated neural calcium activity in normal animals. Dashed blue square indicates the ROI used for calcium trace shown in blue (bottom). Dashed white circle indicates the location of the valve that presses down on *Hydra* when inflated. Gray trace shows the stimulus protocol, where vertical lines indicate valve ‘on’ times. Stimulus applied beginning t = ∼ 20 min and ends t = ∼80 min. Valve is ‘on’ for 1 sec and ‘off’ for 30 sec. (Playback 100x.)

**Video 4:**

Spontaneous ectodermal epitheliomuscular calcium activity in normal animals. Dashed blue square indicates the ROI (entire frame) used for the calcium trace shown in blue (bottom). (Playback 100x.)

**Video 5:**

Stimulated ectodermal epitheliomuscular calcium activity in normal animals. Dashed blue square indicates the ROI (entire frame) used for calcium trace shown in blue (bottom). Dashed white circle indicates the location of the valve that presses down on *Hydra* when inflated. Gray trace shows the stimulus protocol, where vertical lines indicate valve ‘on’ times. Stimulus applied beginning t = ∼ 20 min and ends t = ∼80 min. Valve is ‘on’ for 1 sec and ‘off’ for 30 sec. (Playback 100x.)

**Video 6:**

Spontaneous neural calcium activity in longitudinally bisected animals. Dashed blue circle indicates the ROI used for calcium trace shown in blue (bottom). (Playback 100x.)

**Video 7:**

Stimulated neural calcium activity in longitudinally bisected animals. Dashed blue circle indicates the ROI used for calcium trace shown in blue (bottom). Dashed white circle indicates the location of the valve that presses down on *Hydra* when inflated. Gray trace shows the stimulus protocol, where vertical lines indicate valve ‘on’ times. Stimulus applied beginning t = ∼ 20 min and ends t = ∼80 min. Valve is ‘on’ for 1 sec and ‘off’ for 30 sec. (Playback 100x.)

**Video 8:**

Spontaneous neural calcium activity in headless animals. Dashed blue circle indicates the ROI used for calcium trace shown in blue (bottom). Animal is less active without the head and fewer spikes in calcium activity of peduncle neurons are observed. (Playback 100x.)

**Video 9:**

Stimulated neural calcium activity in headless animals. Dashed blue circle indicates the ROI used for calcium trace shown in blue (bottom). Dashed white circle indicates the location of the valve that presses down on *Hydra* when inflated. Gray trace shows the stimulus protocol, where vertical lines indicate valve ‘on’ times. Stimulus applied beginning t = ∼ 20 min and ends t = ∼80 min. Valve is ‘on’ for 1 sec and ‘off’ for 30 sec. Animal is less active without the head and fewer spikes in calcium activity of peduncle neurons are observed without stimulation. After stimulation begins, more single pulse contractions occur. (Playback 100x.)

**Video 10:**

Spontaneous ectodermal epitheliomuscular calcium activity in footless animals. Dashed blue square indicates the ROI (entire frame) used for the calcium trace shown in blue (bottom). (Playback 100x.)

**Video 11:**

Stimulated ectodermal epitheliomuscular calcium activity in footless animals. Dashed blue square indicates the ROI (entire frame) used for calcium trace shown in blue (bottom). Dashed white circle indicates the location of the valve that presses down on *Hydra* when inflated. Gray trace shows the stimulus protocol, where vertical lines indicate valve ‘on’ times. Stimulus applied beginning t = ∼ 20 min and ends t = ∼80 min. Valve is ‘on’ for 1 sec and ‘off’ for 30 sec. (Playback 100x.)

**Video 12:**

Spontaneous ectodermal epitheliomuscular calcium activity in headless animals. Dashed blue square indicates the ROI (entire frame) used for the calcium trace shown in blue (bottom). Animal is less active without the head and fewer spikes in calcium activity of the whole body ectodermal epitheliomuscular cells are observed. (Playback 100x.)

**Video 13:**

Stimulated ectodermal epitheliomuscular calcium activity in headless animals. Dashed blue square indicates the ROI (entire frame) used for calcium trace shown in blue (bottom). Dashed white circle indicates the location of the valve that presses down on *Hydra* when inflated. Gray trace shows the stimulus protocol, where vertical lines indicate valve ‘on’ times. Stimulus applied beginning t = ∼ 20 min and ends t = ∼80 min. Valve is ‘on’ for 1 sec and ‘off’ for 30 sec. Animal is less active without the head and fewer spikes in calcium activity of epitheliomuscular cells are observed without stimulation. After stimulation begins, more single pulse contractions occur. (Playback 100x.)

**Video 14:**

Spontaneous ectodermal epitheliomuscular calcium activity in bisected animals. Dashed blue square indicates the ROI (entire frame) used for the calcium trace shown in blue (bottom). Animal is less active without the head and fewer spikes in calcium activity of the whole body ectodermal epitheliomuscular cells are observed. (Playback 100x.)

**Video 15:**

Stimulated ectodermal epitheliomuscular calcium activity in bisected animals. Dashed blue square indicates the ROI (entire frame) used for calcium trace shown in blue (bottom). Dashed white circle indicates the location of the valve that presses down on *Hydra* when inflated. Gray trace shows the stimulus protocol, where vertical lines indicate valve ‘on’ times. Stimulus applied beginning t = ∼ 20 min and ends t = ∼80 min. Valve is ‘on’ for 1 sec and ‘off’ for 30 sec. Animal is less active without the head and fewer spikes in calcium activity of epitheliomuscular cells are observed without stimulation. After stimulation begins, more single pulse contractions occur. (Playback 100x.)

**Video 16:**

Spontaneous ectodermal epitheliomuscular calcium activity in body column animals. Dashed blue square indicates the ROI (entire frame) used for the calcium trace shown in blue (bottom). Animal is less active without the head and fewer spikes in calcium activity of the whole body ectodermal epitheliomuscular cells are observed. (Playback 100x.)

**Video 17:**

Stimulated ectodermal epitheliomuscular calcium activity in body column animals. Dashed blue square indicates the ROI (entire frame) used for calcium trace shown in blue (bottom). Dashed white circle indicates the location of the valve that presses down on *Hydra* when inflated. Gray trace shows the stimulus protocol, where vertical lines indicate valve ‘on’ times. Stimulus applied beginning t = ∼ 20 min and ends t = ∼80 min. Valve is ‘on’ for 1 sec and ‘off’ for 30 sec. Animal is less active without the head and fewer spikes in calcium activity of epitheliomuscular cells are observed without stimulation. After stimulation begins, more single pulse contractions occur. (Playback 100x.)

**Video 18:**

Stimulated neural calcium activity in the hypostome and body column of normal animals. Dashed blue circle indicates the ROI used for calcium trace shown in blue (bottom). Dashed white circle indicates the location of the valve that presses down on *Hydra* when inflated. Gray trace shows the stimulus protocol, where vertical lines indicate valve ‘on’ times. Stimulus applied beginning t = ∼ 20 min, Valve is ‘on’ for 1 sec and ‘off’ for 30 sec. Imaged with 10x 0.45 N.A. objective (Playback 100x.)

## Source data

**Figure 2 – source data 1: source file for stimulus dependent mechanosensory response.**

The file (Fig2_SourceData.mat) contains all source data used for quantitatively characterizing mechanosensory response shown in Figure 2. The file contains a struct named ‘HydraData’ with each trial in a different row. The columns contain the raw and processed data for each trial: ‘StimCondition’ indicates the stimulation pressure (0, 5, 10, 15, 20 or 25 psi) used during the trial; ‘RawFluorescenceFoot’ contains the raw fluorescence values from foot/peduncle ROI; ‘StimulationTrace’ contains the trace of stimulus delivery where 0 is valve off, 1 is valve on (and stimulation is being applied) per frame; ‘Response Probability’ is extracted after processing ‘RawFluorescenceFoot’ with stimulation onset times from ‘StimulationTrace’; ‘TimeBetweenContractions’ is time in seconds between contraction burst pulses extracted from ‘RawFluorescenceFoot’; ‘SinglePulseContractions_Percent’ is the percent of body contractions that occur as single pulses extracted from ‘RawFluorescenceFoot’

**Figure 3 – source data 1: source file mechanosensory response in resected animals.**

The file (Fig3_SourceData.mat) contains all source data used for characterizing mechanosensory response in resected animals in Figure 3. The file contains a struct named ‘HydraData’ with each trial in a different row. The columns contain the raw and processed data for each trial: ‘StimCondition’ indicates stimulation condition (w/ or w/o stimulation; body resections - whole, bisected, no foot = Footless, no head = Headless, no head and foot = BodyColumn); ‘RawFluorescenceFoot’ contains the raw fluorescence values from foot/peduncle ROI; ‘StimulationTrace’ contains the trace of stimulus delivery where 0 is valve off, 1 is valve on (and stimulation is being applied) per frame; ‘Response Probability’ is the stimulated contraction probability extracted after processing ‘RawFluorescenceFoot’ with stimulation onset times from ‘StimulationTrace’; ‘BasalContractionProbability’ is spontaneous contraction probability extracted after processing ‘RawFluorescenceFoot’ during (t = 0-20 min) the initial acclimation period without stimulation.

**Figure 4 – source data 1: source file for fluorescent calcium activity from single neuron ROIs.**

The file (Fig4_SourceData.zip) contains three folders ‘Hydra1’, ‘Hydra2’ and ‘Hydra3’ for the three different animals with individually tracked single neurons. Each folder contains a file called ‘IndividualNeuronTraces.mat’ with three different matrices: ‘NeuronROI_RawFluorAllFrames’ matrix [# of rows = # of neurons+1, # of columns = # of frames] contain the raw fluorescence intensities individual neuronal ROIs. Each row is different neuron and the last row in the matrix is the average fluorescence from peduncle ROI’; ‘ROI_xLocAllFrames’ matrix [# of rows = # of neurons, # of columns = # of frames] contains the x position of the center of circular ROIs for individual neurons; ‘ROI_yLocAllFrames’ matrix [# of rows = # of neurons, # of columns = # of frames] contains the y position of the center of circular ROIs for individual neurons

