## Supplementary figures for "Multiple neuronal networks coordinate *Hydra* mechanosensory behavior"

### Supplementary Information

### Supplementary Figures:

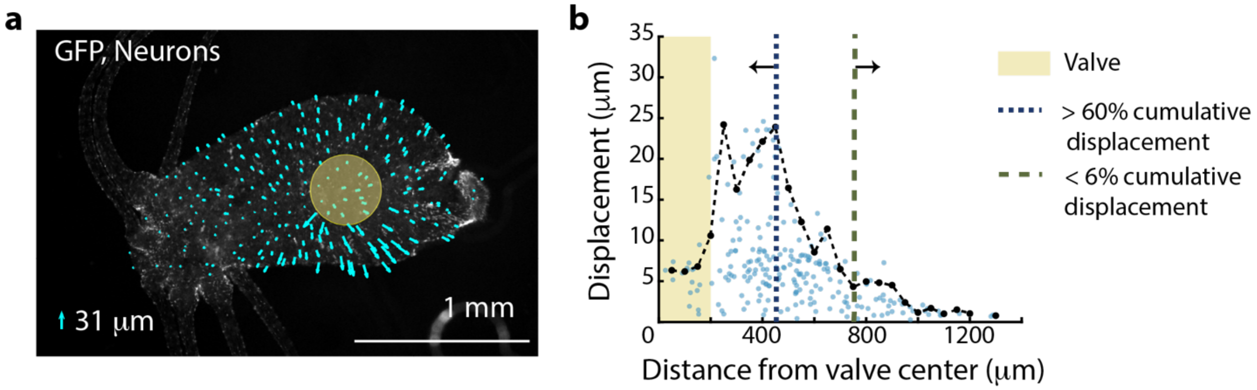


**Figure 2 – Figure Supplement 1: Distribution of mechanical forces.** Fluorescent image (grayscale) of transgenic *Hydra* (nGreen) expressing GFP in neurons and neural progenitors. The Oral region is on the left. Aboral//peduncle region is on the right. Arrows (cyan) overlayed indicate the displacement of individual cells immediately before and after the valve is pressurized for stimulation (~50 msec). The lengths of the arrows are enlarged to be visible with the scaled size of the largest displacement ~31 µm as indicated by the legend (lower left). Semi-transparent (yellow) circle indicates the location of the valve (400 µm diameter). b) Displacement of neurons due to stimulation varies with the location of the neurons from the valve center. Each semi-transparent blue dot represents a single neuron (n= 222 neurons). Black dots (connected by black dashed line) represent the average displacement of the highest three displacements within a 50 µm radial band. Microfluidic valve has a radius of 200 µm indicated by the yellow shaded region. Majority of the cumulative displacement (>60 %) is localized to a 250 µm radius bordering the valve indicated by the blue dashed line at 450 µm. Neurons more than 750 µm from the microfluidic valve center have negligible displacement (<6% cumulative displacement) indicated by the green dashed line.

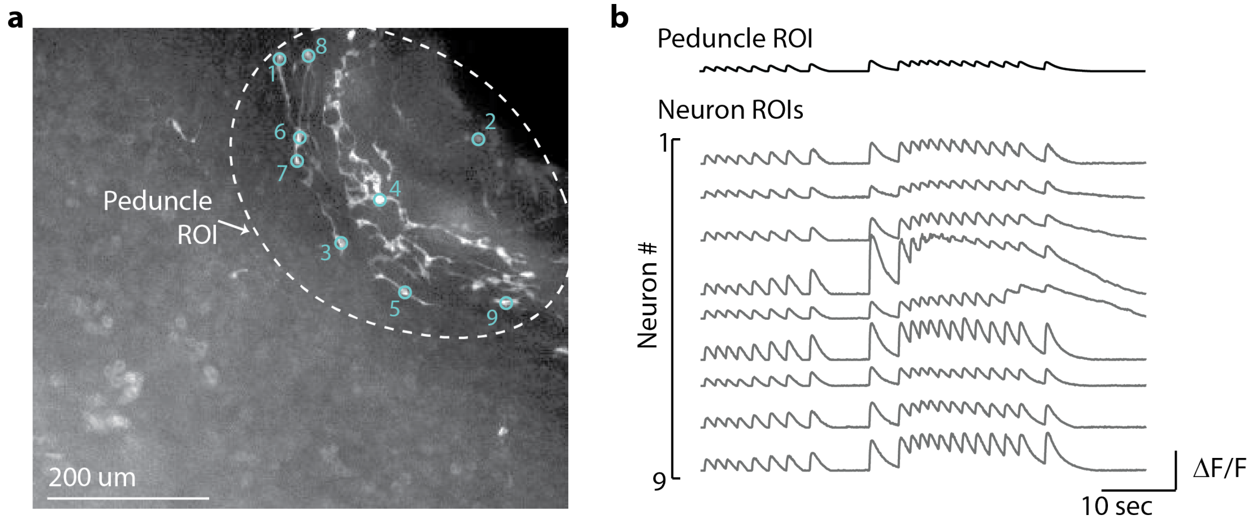


**Figure 2 – Figure Supplement 2: Average calcium fluorescence from large peduncle ROI correlated with calcium fluorescence from smaller ROIs for individual peduncle neurons.** a) Fluorescent image of transgenic *Hydra* expressing GCaMP6s pan-neuronally. Dashed while circle indicates the peduncle ROI. Small cyan circles indicate the individual neuron ROIs. Peduncle region is in the top right. The oral end of the *Hydra* (not shown in the image) is oriented towards the bottom left corner. b) Average calcium fluorescence trace (black) from the large peduncle ROI. Average calcium traces (gray) from individual neurons in the peduncle.

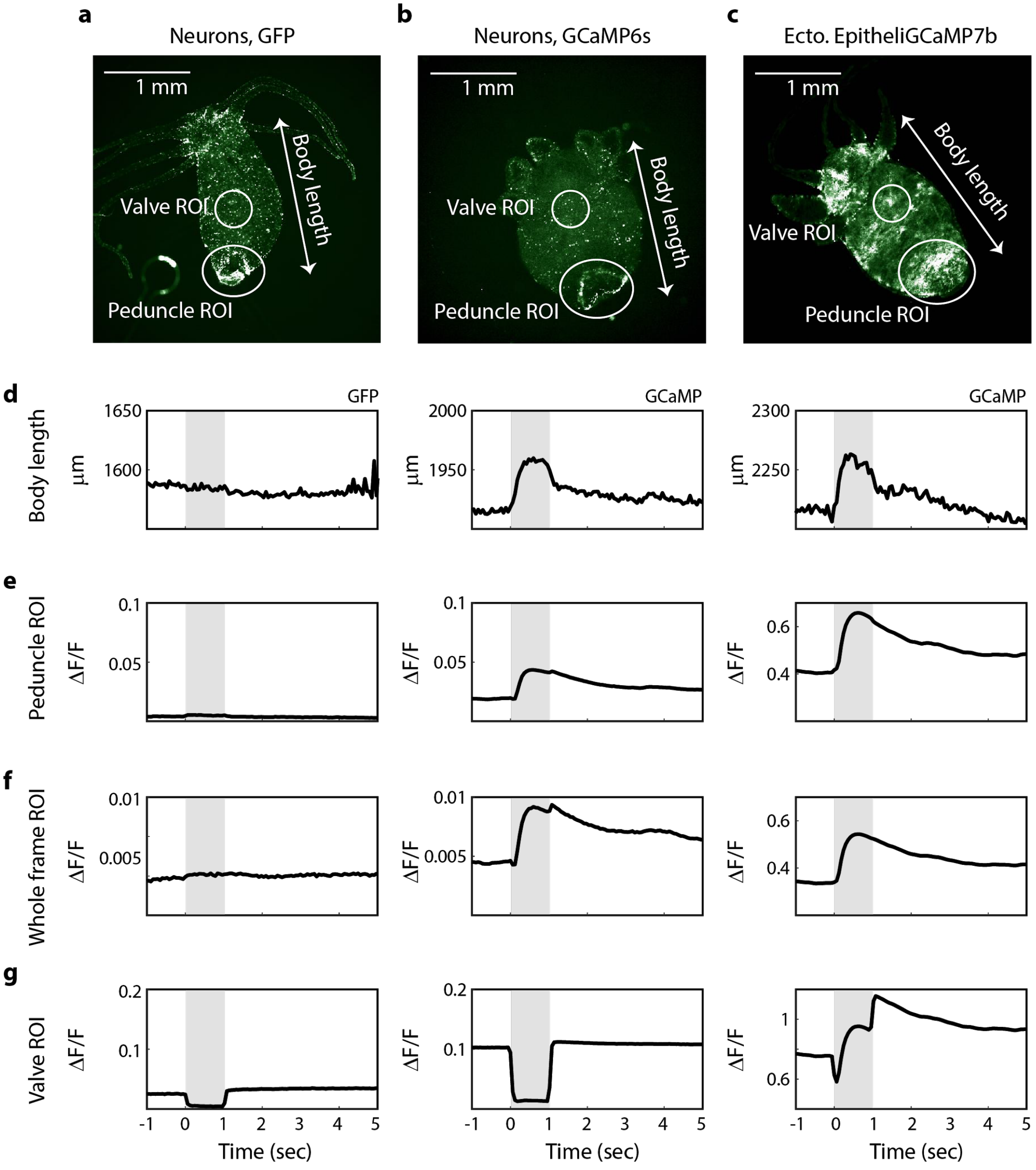


**Figure 2 – Figure Supplement 3: Spikes in calcium fluorescence are due to calcium activity not motion artifacts.** a) Fluorescence image of transgenic *Hydra* (nGreen) expressing GFP in neurons false colored with hot green colormap. Neurons appear in white. b) Fluorescence image of transgenic *Hydra* expressing GCaMP6s in neurons false colored with hot green colormap. c) Fluorescence image of transgenic *Hydra* expressing GCaMP7b in ectodermal epitheliomuscular cells false colored with hot green colormap. a-c) Body length (white arrow), valve ROI (top circles), peduncle ROI (larger bottom circles) and whole frame ROI (entire frame square) annotated on all three images for comparison. d) Average body length trace time-aligned to stimulus onset from GFP *Hydra* (left) and GCaMP *Hydra* (middle, right). Light gray shaded region indicates when the stimulation is on. e) Average fluorescence trace calculated from the annotated peduncle ROI from GFP *Hydra* (left) and GCaMP *Hydra* (middle, right). f) Average fluorescence trace calculated from the whole frame ROI from GFP *Hydra* (left) and GCaMP *Hydra* (middle, right). g) Average fluorescence trace calculated from the annotated valve ROI from GFP *Hydra* (left) and GCaMP *Hydra* (middle, right).

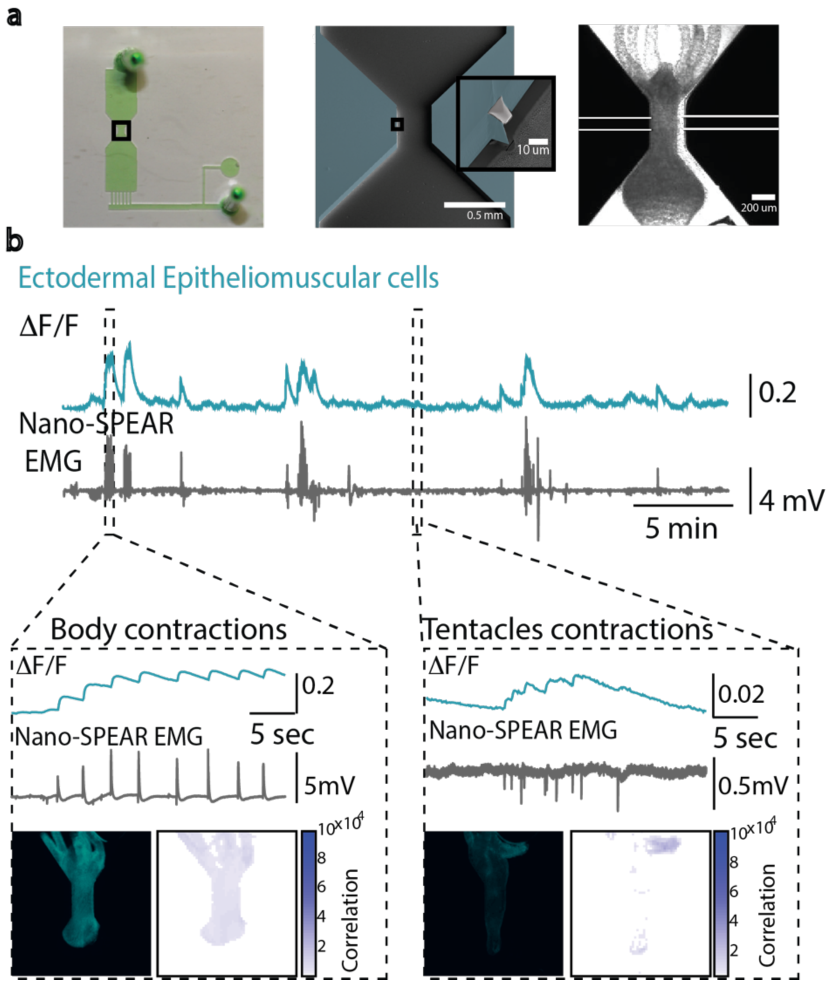


**Figure 2 – Figure Supplement 4: Simultaneous electrophysiology and calcium imaging of ectodermal epitheliomuscular cells.** (a) Photograph (left) of a microfluidic immobilization chamber filled with green dye. Black box highlights the recording region in the microfluidic chamber (110 μm tall). False colored scanning electron micrograph (middle) shows the recording region (blue, photoresist; light grey, Pt; dark grey, silica) on the nano-SPEAR chip (50 μm tall). Inset shows a zoom-in of the Pt electrode (light grey) suspended mid-way between the top and bottom of the photoresist sidewall (blue). Brightfield image (right) shows Hydra immobilized in the microfluidic chamber placed on top of the nano-SPEAR chip with combined 160 μm tall recording region (Reproduced from Figure 2A, Badhiwala et al, 2018[31] by permission of The Royal Society of Chemistry. This panel is not covered by the CC-BY 4.0 license, and further reproduction of this panel would need permission from the copyright holder.) b) Simultaneous electrophysiology and calcium imaging in transgenic *Hydra* (GCaMP6s, ectodermal epitheliomuscular cells). Top trace shows mean fluorescence ($\Delta F/F$) from ectodermal epitheliomuscular cells (whole frame ROI) in *Hydra* (GCaMP, ectodermal) (10Hz). Bottom trace shows simultaneously recorded electrical activity from the *Hydra* (1KHz). High and low activity periods identified based on peak amplitude. Inset: left box shows correlation during high activity period, contraction bursts. Traces show peaks in fluorescence (top trace) coinciding with peaks in electrophysiology (bottom trace). A representative fluorescence image shows high levels of fluorescence thus calcium activity in the entire body during contraction burst events. Correlation map spatially plotting the correlation coefficient shows the entire body has calcium activity correlated with electrophysiology. Inset: right box shows correlation during low activity period, tentacle pulses. Traces show peaks in fluorescence (top trace) coinciding with peaks in electrophysiology (bottom trace). A representative fluorescence image shows high levels of fluorescence in the tentacle region during tentacle pulses. Cross correlation map shows the tentacle region has calcium activity correlated with electrophysiology. Almost all spikes in electrophysiology coincide with spike in fluorescence from ectodermal epitheliomuscular cells (and behavioral contractions).

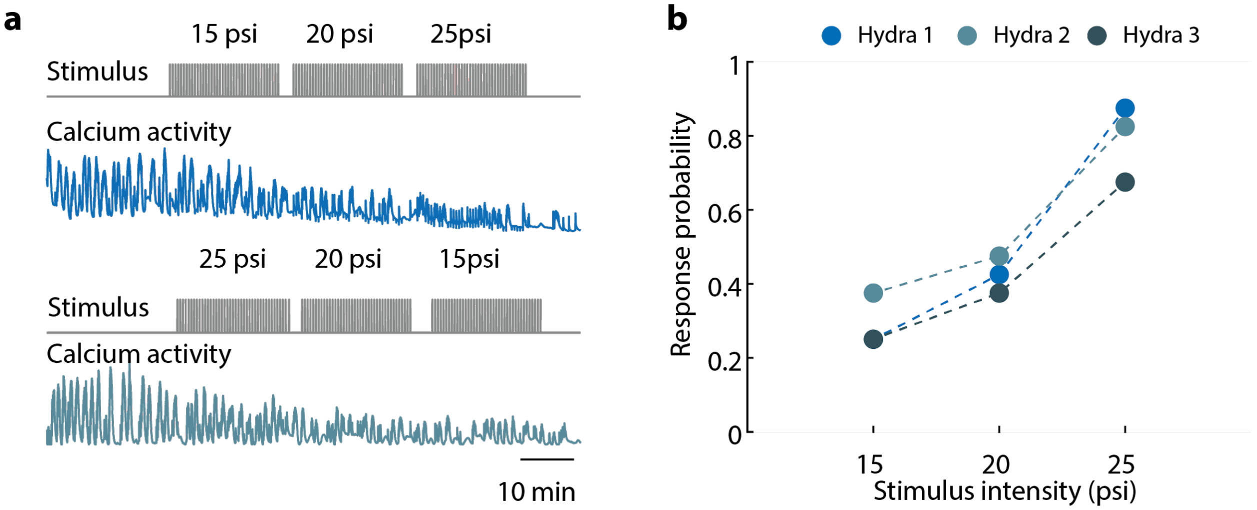


**Figure 2 – Figure Supplement 5: *Hydra*'s epitheliomuscular response is dependent on the mechanical stimulus intensity**. a) Gray trace (top) is the stimulation protocol. 20 min no stimulation, 20 min of repeated stimulation (1 sec 'on', 30 sec 'off') at 15 psi, 20 min of repeated stimulation (1 sec 'on', 30 sec 'off') at 20 psi, and 20 min of repeated stimulation (1 sec 'on', 30 sec 'off') at 25 psi. Stimulus 'on' times indicated by vertical lines. Entire frame ROI used for analysis of the whole-body epithelial calcium activity. Representative calcium fluorescence trace (blue) from ectodermal epitheliomuscles (GCaMP7b) from animal stimulated with three different stimulus intensities for 20 min each (15, 20 and 25 psi). Representative calcium fluorescence trace (teal) from ectodermal epitheliomuscles (GCaMP7b) from animal stimulated with three different stimulus intensities for 20 min each (25, 20 and 15 psi). The decrease in fluorescence amplitude observed for both increasing in and decreasing stimulus intensity is due to photobleaching of the calcium indicator. b) Mechanosensory response probability at different stimulus intensities (N = 3 animals). Response probability, fraction of trials that have at least one calcium spike (also contraction pulse) occurring within 1 sec of stimulation onset.

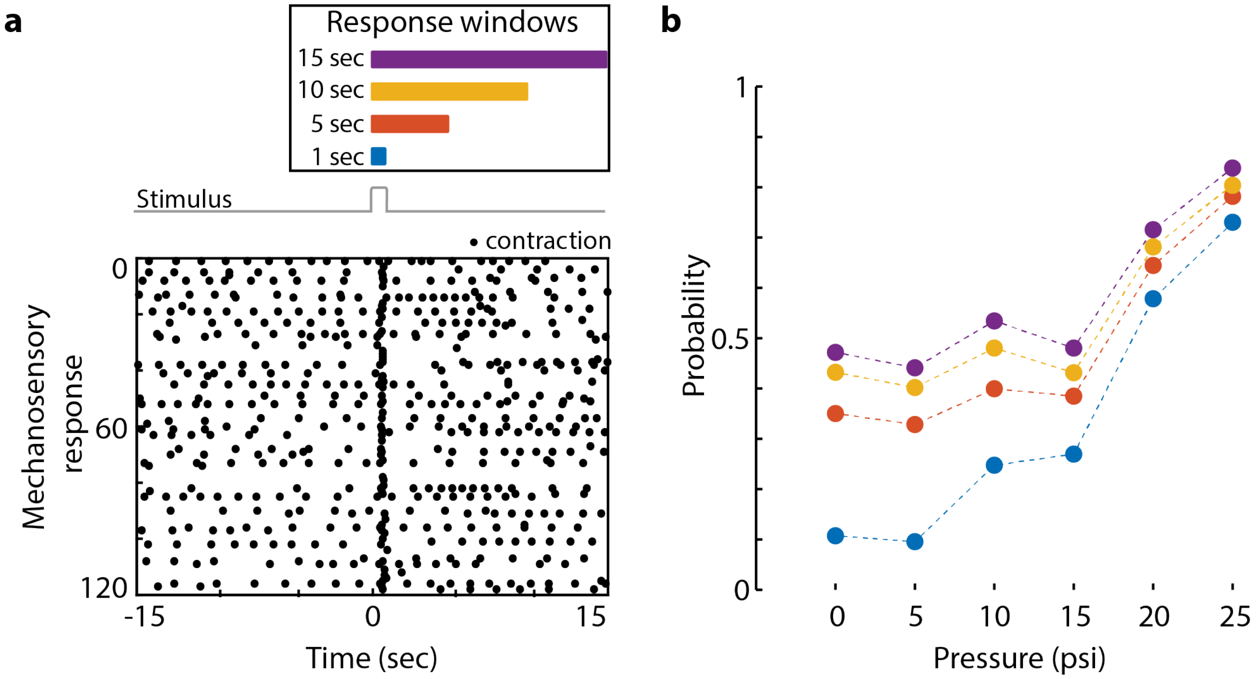


**Figure 2 – Figure Supplement 6: Mechanosensory response window.** a) Different response windows tested (1 sec in blue; 5 sec in orange; 10 sec in yellow; and 15 sec in purple) (Top). The lengths of the rectangles correspond to time (sec) in the raster plot below. Stimulus trace in gray. Representative raster plot of time-aligned contraction pulses from multiple trials superimposed from one animal stimulated at 20 psi every 31 sec for 60 min (Bottom). Each black dot is a spike in calcium fluorescence identified as a contraction pulse. Stimulus is applied from 0 - 1 sec as indicated by a step in stimulus trace. b) Response probability, fraction of trials that have at least one calcium spike (also contraction pulse) occurring during different response windows (1 sec in blue; 5 sec in orange; 10 sec in yellow; and 15 sec in purple within stimulation onset).

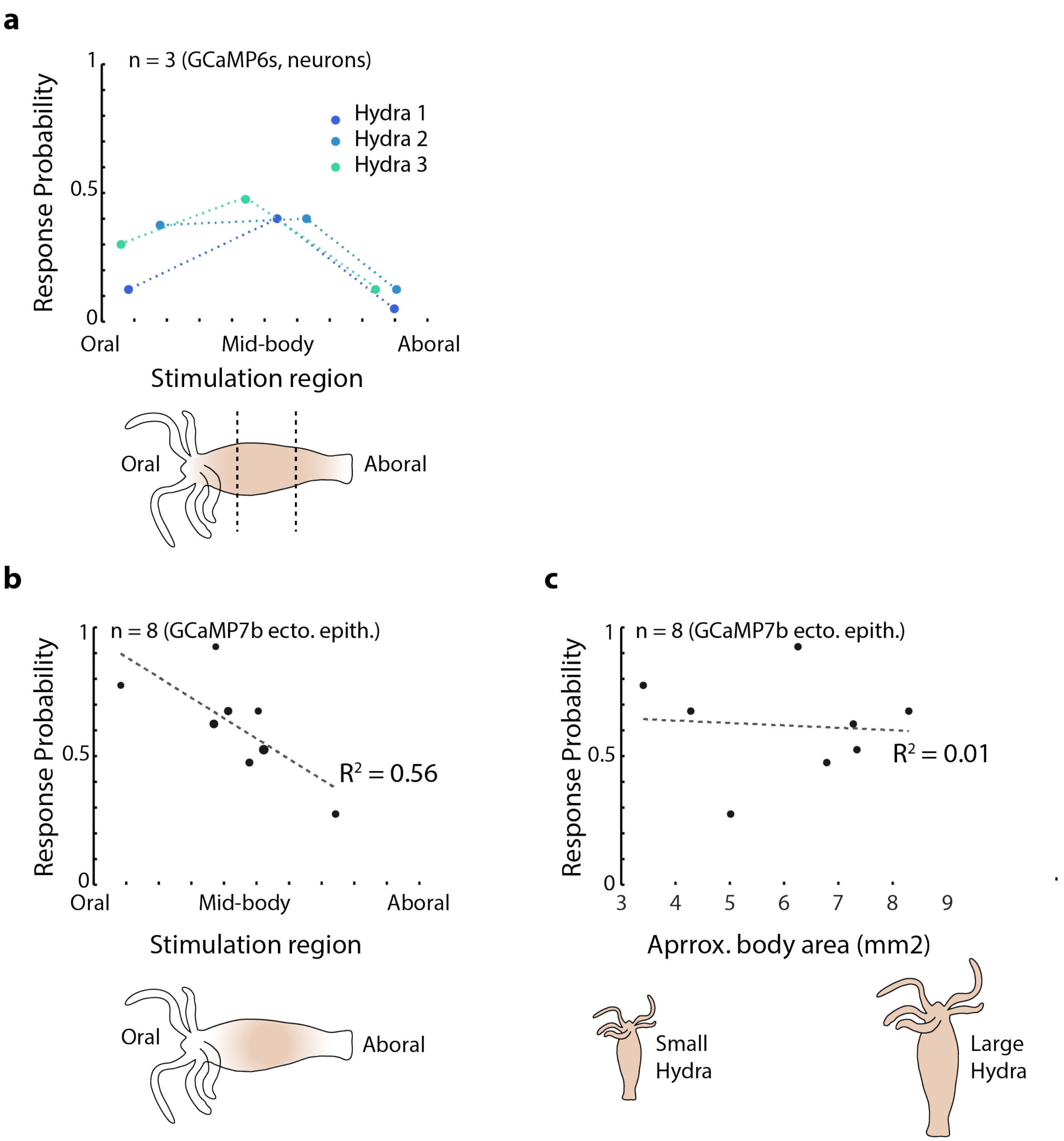


**Figure 2 – Figure Supplement 7: Mechanical sensitivity of different body regions in *Hydra*.** a) Response probability of transgenic Hydra (N=3 animals expressing GCaMP6s in neurons) stimulated at three different body regions: Oral, Mid-body and Aboral. Annotated Hydra below the plot indicates the three stimulation regions used. b) Response probability of transgenic Hydra (n=8 animal expressing GCaMP7b in ectodermal epitheliomuscular cells) Response probability is calculated using average calcium fluorescence from neurons in the peduncle ROI in a) and using average calcium fluorescence from the ectodermal epitheliomuscular cells from the entire body in b) and c).

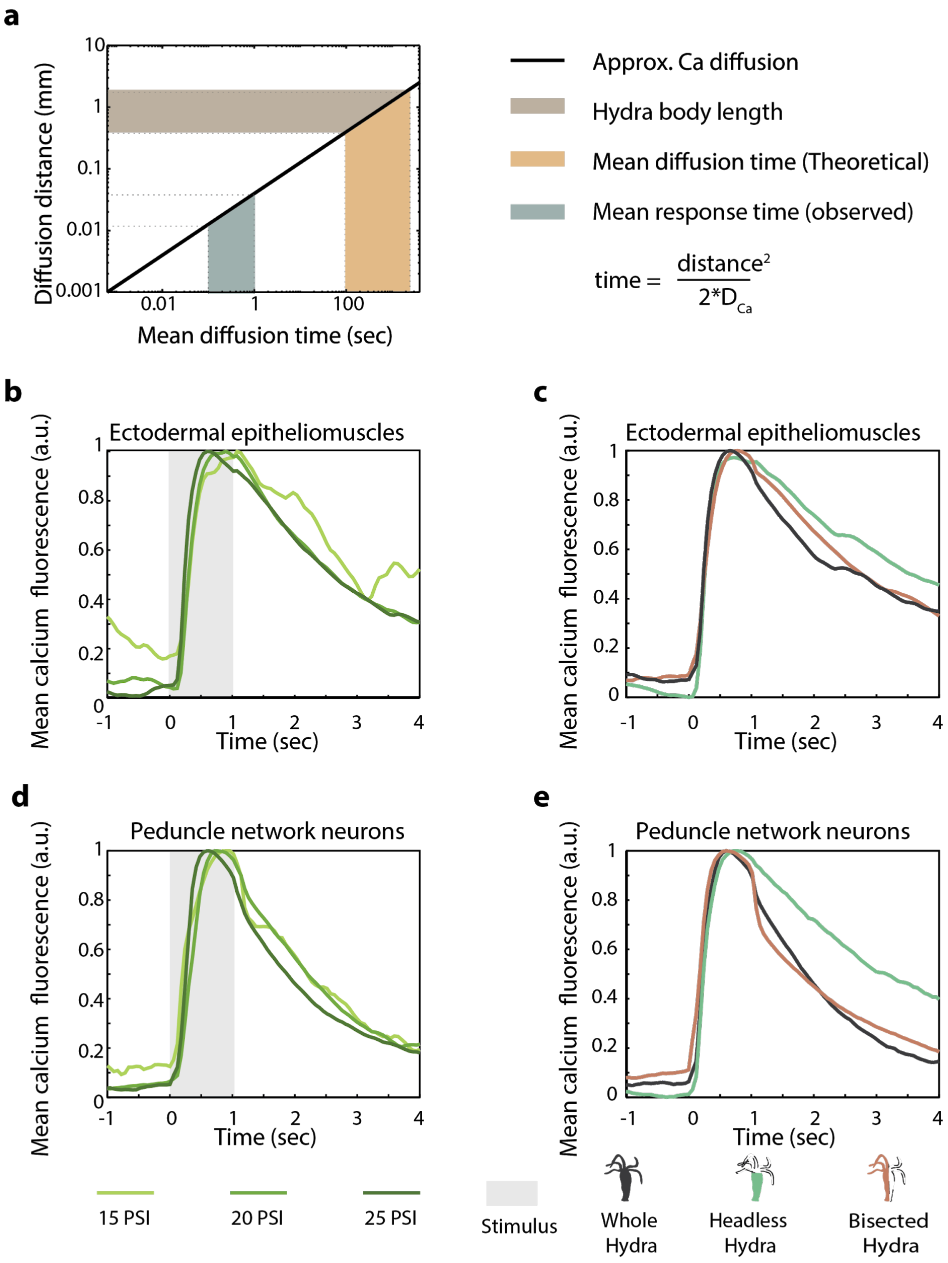


**Figure 2 – Figure Supplement 8: *Hydra's* mechanosensory response time is faster than passive calcium diffusion through epitheliomuscular cells** a) Distance calcium diffuses passively over given time (left, black line plot) approximated using passive diffusion equation (right). Light brown shaded region indicates the range of body length (~ 0.5 - 1 mm) of *Hydra* in microfluidic chambers used for the experiments. Teal shaded region indicates the average time between stimulus onset and observed spike in fluorescence (mean response time 0.5 - 1 sec). Yellow shaded region is the theoretical mean calcium diffusion time calculated assuming passive diffusion. b-e) Mean calcium fluorescence time-aligned to stimulus and normalized. Stimulus is applied at 0 sec (valve ‘on’ from 0 - 1 sec). b) Mean calcium activity from the peduncle network of neurons at different stimulus intensities. c) Mean calcium activity from ectodermal epitheliomuscular cells at different stimulus intensities. b-c) The line colors correspond to different stimulus intensities shown at the bottom. d) Mean calcium activity from peduncle network of neurons in different body resections stimulated at 20 psi (pressure for ~ 60% response probability in whole animals). e) Mean calcium activity from ectodermal epitheliomuscular cells in different body resections different body resections stimulated at 22 psi (pressure for ~ 60% response probability in whole animals). d-e) The line colors correspond to the different resections shown at the bottom with cartoon schematic of the resected *Hydra* with the same color.

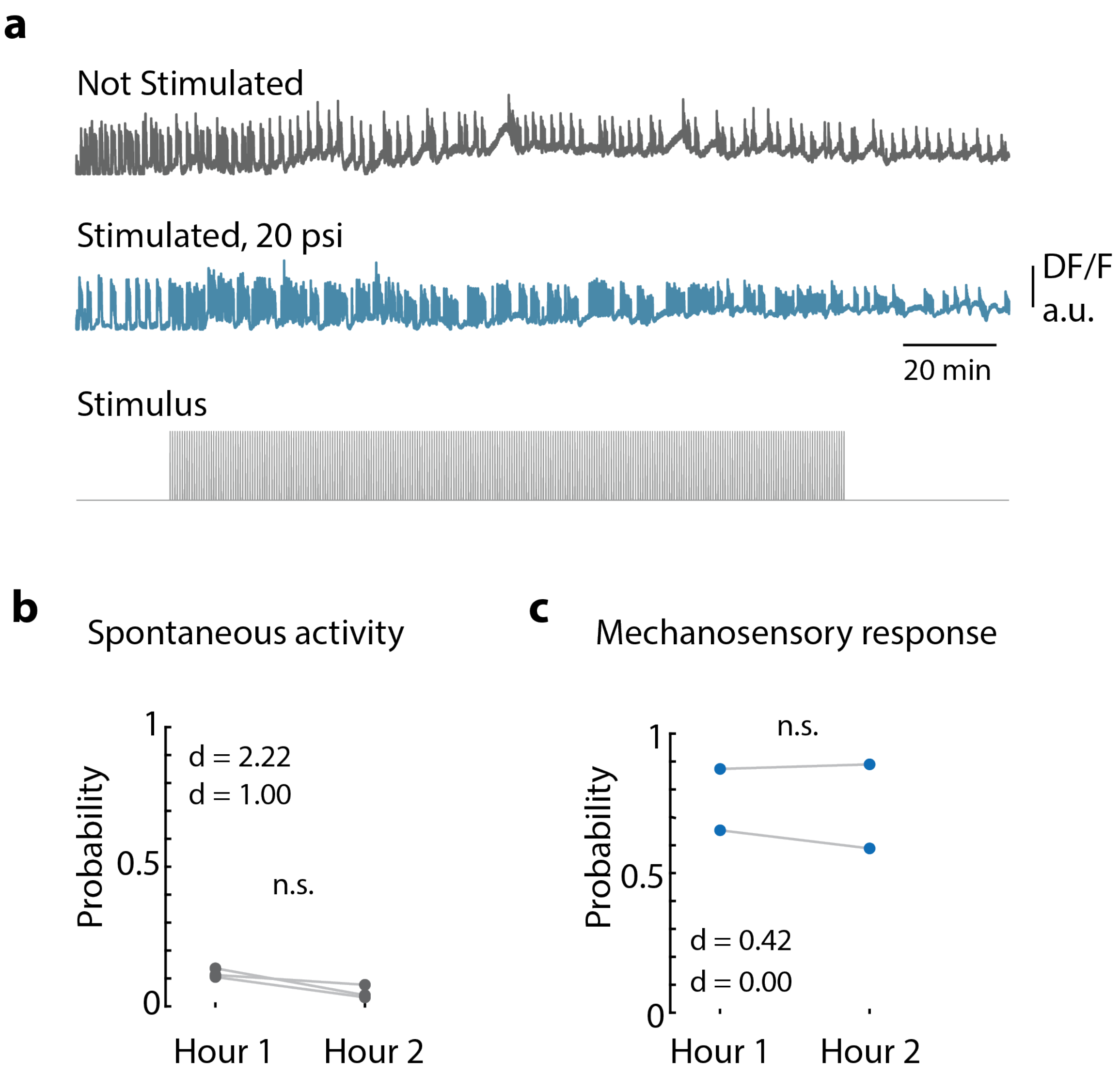


**Figure 2 – Figure Supplement 9: Long-term mechanical stimulation** a) Representative calcium fluorescence trace from the peduncle region from an animal (GcaMP6s, neurons) not stimulated and animal mechanically stimulated with 20 psi. Stimulus protocol in gray, 20 min no stimulation, 120 min of repeated stimulation (1 sec 'on', 30 sec 'off') at 20 psi followed by no stimulation. b) Spontaneous and stimulated (mechanosensory response) contraction pulse probabilities for the first and second hour of stimulation compared for each animal. Gray circles are mean contraction pulse probability during no stimulation calculated from an average of 1 sec window shifted by ~ 0.3 sec over 30 sec intervals (N = 3). Blue circles are contraction probability during stimulation calculated from 1 sec response window during valve ‘on’ (N = 2). Light gray lines indicate the change in probabilities from hour 1 to hour 2 for each individual. (paired t-test, n.s. = not significant)

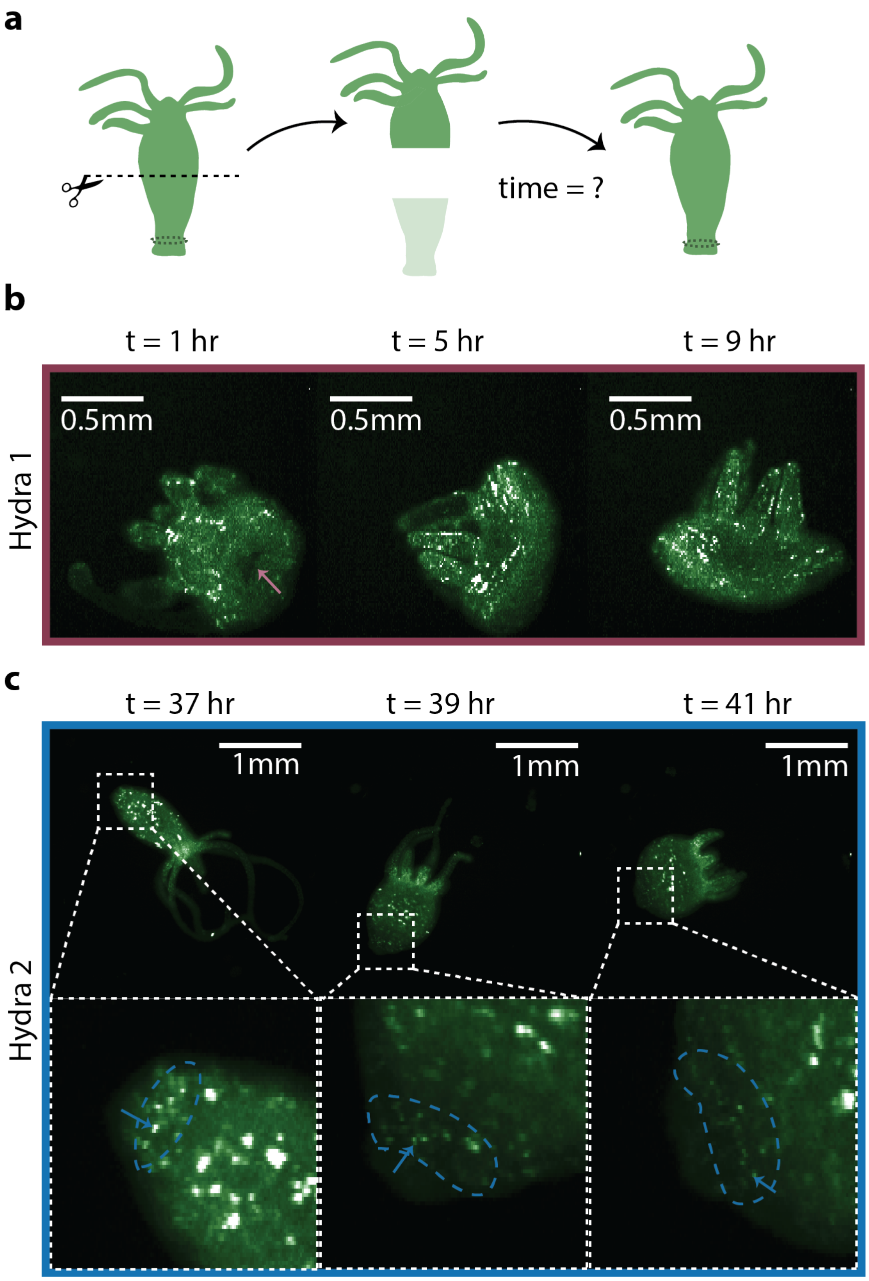


**Figure 3 – Figure Supplement 1: Regeneration of the peduncle network.** a) Summary schematic of the experiment performed. Dashed line indicates where the incision was made to remove the lower half of the transgenic *Hydra* body (GcaMP6s, neurons) to discover when the peduncle neuronal network is regenerated. b) Fluorescence images of "footless" *Hydra* during body contractions at various timepoints t = 1 hr, 5 hr and 9 hr after bisection. Pink arrow indicates the open wound visible at t = 1 hr but not at other timepoints. c) Fluorescence images of another "footless" *Hydra* during body contractions at various timepoints t = 37 hr, 39 hr, 41 hr. Blue dashed circle indicates the region where the peduncle network is found. Blue arrows indicate some of the neurons in the peduncle network with clearly well-connected neurons at t = 41 hr.

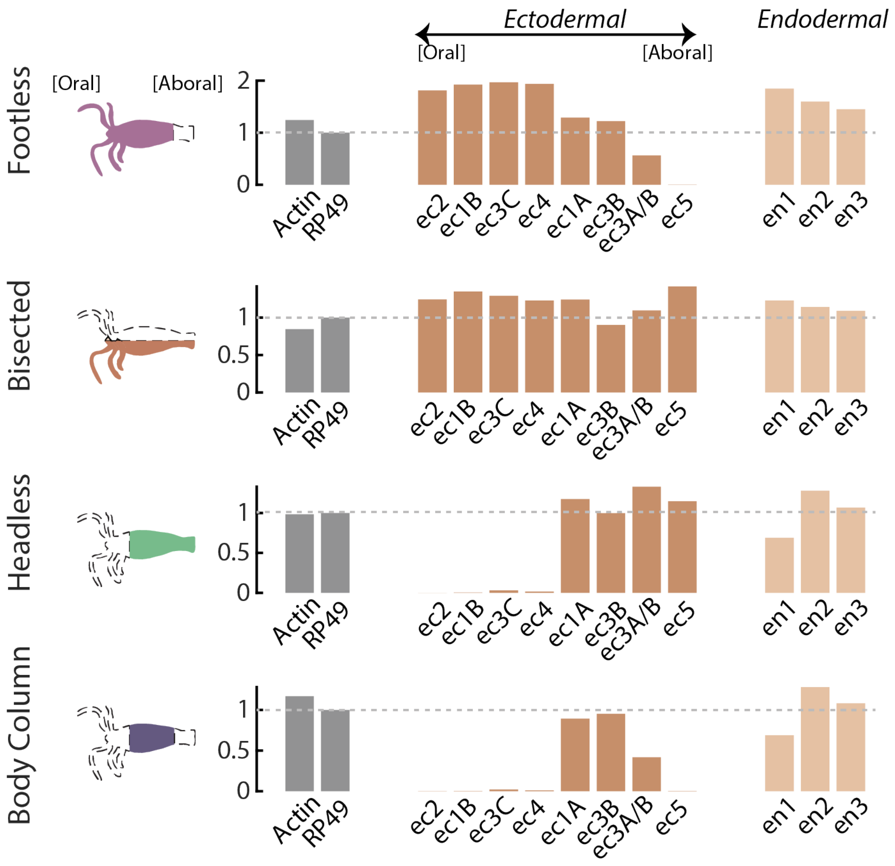


**Figure 3 - Figure Supplement 2: RT-qPCR analyses of neuron subtype-specific gene expression in resected animals demonstrates loss of specific neuron subtypes.** RT-qPCR was used to test for the loss of specific neuron subtypes in whole, tube (no head or foot), headless, footless, and bisected *Hydra* using uniquely expressed biomarkers for each subtype. There is not a specific biomarker for ec3A (located in the basal disk), so the marker used to test for the presence or absence of this cell type is also expressed in ec3B (located in the body column). Therefore, expression of the ec3A/B marker gene is reduced, but not completely lost in animals with resected feet ("tube" and "footless"). However, ec5 expression (located in the peduncle above the basal disk) is completely lost in animals with resected feet, thus it is clear that the ec3A subtype is completely lost in these animals. Biomarkers for neuron subtypes located in the head and tentacles (ec1B, ec2, ec3C, ec4) are completely lost in animals with resected heads (“headless” and “tube”). Data were analyzed with the $2^{-\Delta\Delta Ct}$ method and results were normalized to the housekeeping gene *RP49* and to expression in whole animals.

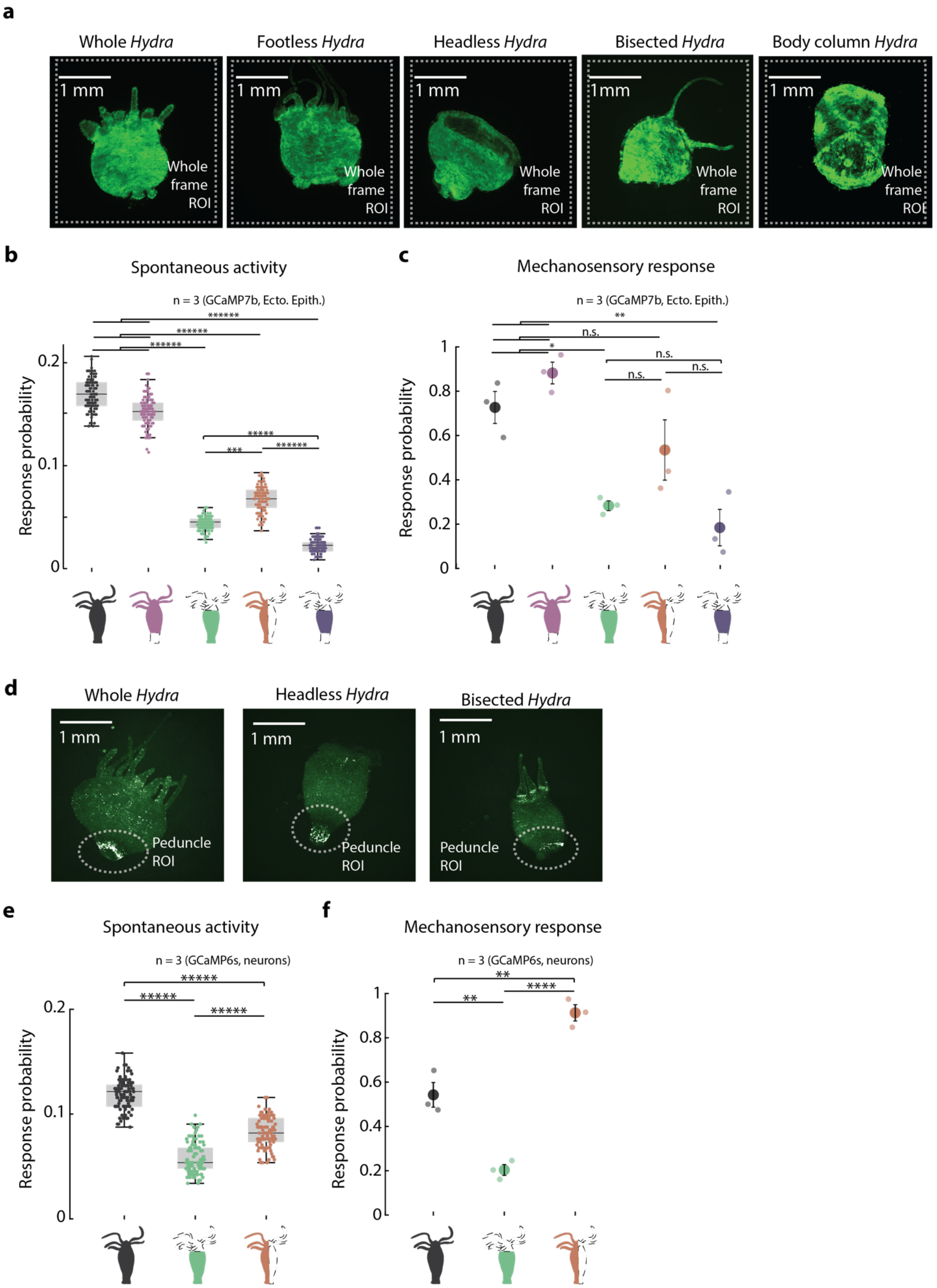
a
Whole Hydra
Footless Hydra
Headless Hydra
Bisected Hydra
Body column Hydra
1 mm
Whole frame ROI
b
Spontaneous activity
n = 3 (GCaMP7b, Ecto. Epith.)
Response probability
c
Mechanosensory response
n = 3 (GCaMP7b, Ecto. Epith.)
n.s.
d
Whole Hydra
Headless Hydra
Bisected Hydra
Peduncle ROI
e
Spontaneous activity
n = 3 (GCaMP6s, neurons)
f
Mechanosensory response
n = 3 (GCaMP6s, neurons)

**Figure 3 – Figure Supplement 3: Mechanosensory response from ectodermal epitheliomuscular cells and neurons in resected *Hydra*.** a) Representative images of resection preparations of transgenic *Hydra* (GCaMP7b, ectodermal epitheliomuscular cells: whole (or control), “footless”, “headless”, “bisected”, and “body column”. White dashed square indicates the whole frame ROI used for quantifying response. b) Spontaneous probability of at least one spiking event (or contraction pulse) occurring during a random 1 sec window. (1 sec window slide by ~ 0.3 sec across 30 sec stimulation intervals; Kruskal-Wallis test with post hoc dunn-sidak correction) c) Mechanosensory response probability during 1 sec of valve on (Response probability, fraction of trials that have at least one calcium spike (also contraction pulse) occurring within 1 sec of stimulation onset). Large circles indicate average values from all animals combined for each condition. Small circles indicate probability from individual animals (one-way anova with post hoc Bonferroni correction). Error bars are standard error of mean (S.E.M.) (N = 3 *Hydra* for each condition; analysis from 60 min of stimulation which is imaging t = 20 – 80 min; n.s. = not significant, ** = $p < 0.01$, **** = $p < 0.0001$, ***** = $p < 0.00001$). d) Representative images of resection preparations of transgenic *Hydra* (GCaMP6s, neurons: whole (or control), “headless”, and “bisected”. White dashed circle indicates the peduncle ROI used for quantifying response. e) Spontaneous probability of at least one spiking event (or contraction pulse) occurring during a random 1 sec window. (1 sec window slide by ~ 0.3 sec across 30 sec stimulation intervals; Kruskal-Wallis test with post hoc dunn-sidak correction) f) Mechanosensory response probability during 1 sec of valve on. Large circles indicate average values from all animals combined for each condition. (Response probability, fraction of trials that have at least one calcium spike (also contraction pulse) occurring within 1 sec of stimulation onset.) Small circles indicate probability from individual animals (one-way anova with post hoc Bonferroni correction). Error bars are standard error of mean (S.E.M.) (N = 3 *Hydra* for each condition; analysis from 60 min of stimulation which is imaging t = 20 – 80 min; n.s. = not significant, ** = $p < 0.01$, **** = $p < 0.0001$, ***** = $p < 0.00001$).

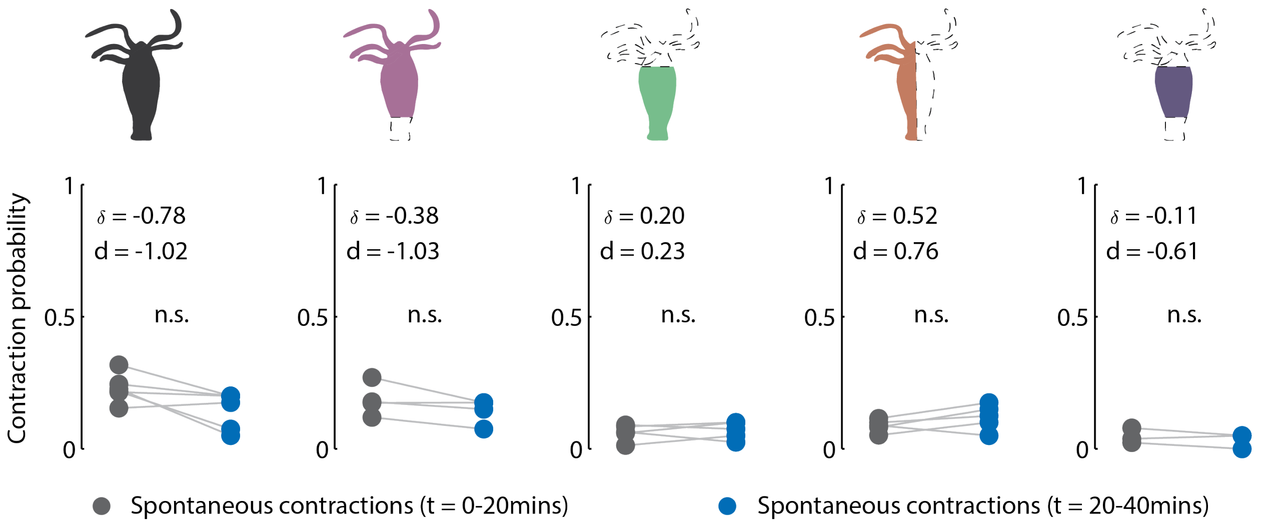


**Figure 3 – Figure Supplement 4: Contraction activity in non-stimulated animals.** Cartoon schematic of resection preparations of transgenic *Hydra* (GCaMP7b, ectodermal epitheliomuscular cells): whole (or control), “footless”, “headless”, “bisected” and “body column” animals. Entire frame ROI used for analysis of the whole-body epithelial calcium activity. Gray circles are mean contraction probability during t = 0 - 20 min. Blue circles are mean contraction probability during t = 20 - 40 min, (this time corresponds to when stimulated animals receive mechanical stimuli). Mean probability calculated from 1 sec window shifted by ~ 0.3 sec over 30 sec intervals. (Response probability, fraction of trials that have at least one calcium spike (also contraction pulse) occurring within 1 sec of stimulation onset). Pressure in valves = 0 psi. Light gray lines pair the spontaneous contractions probability from the first 20 min interval with the second 20 min interval from each individual animal. (Whole N = 4, footless N = 3, headless N = 3, bisected N = 5, body column N = 3 *Hydra*, paired t-test, n.s. = not significant)

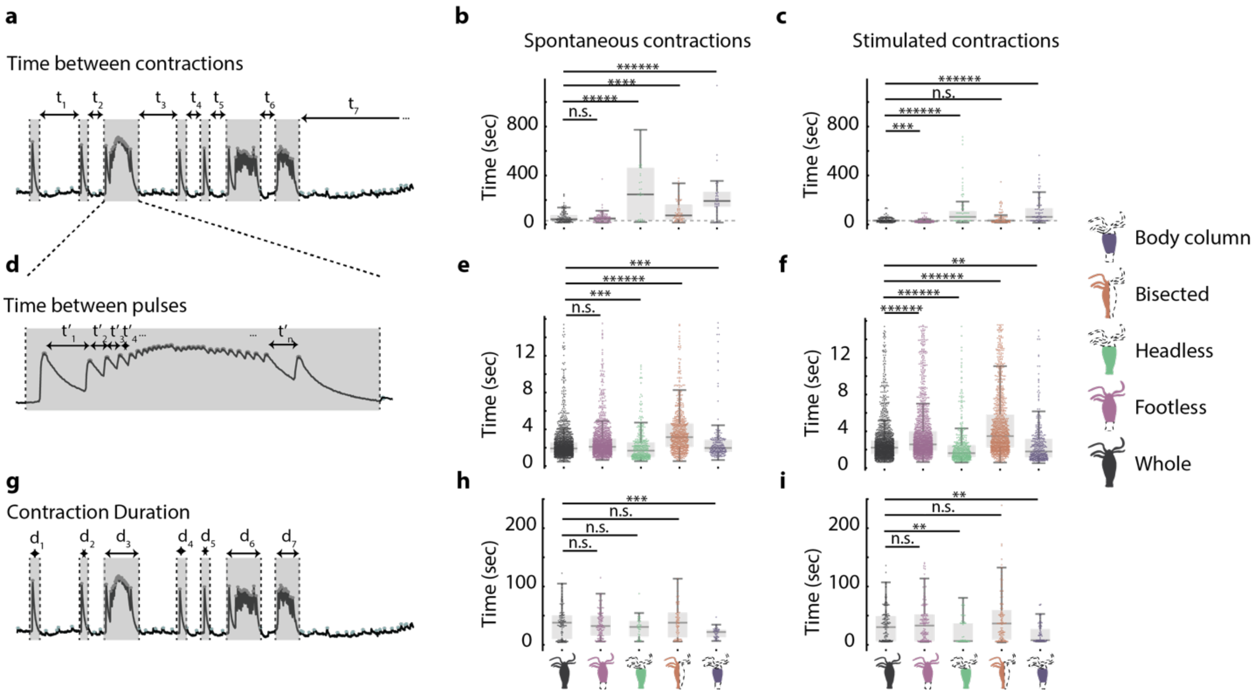


**Figure 3 – Figure Supplement 5: Hypostome and peduncle nerve rings work together to coordinate contractile behavior** a) Representative fluorescence trace used to calculate time interval between contractions. b) Time interval between spontaneous contractions in animals with different resections: whole, “footless”, “headless”, “bisected” and “body column”. c) Time interval between stimulated contractions in animals with different resections. d) Representative fluorescence trace used to calculate time interval between contraction pulses. For illustration purpose, only a select few of the time intervals are shown. e) Time interval between spontaneous contraction pulses in animals with different resections. f) Time interval between stimulated contraction pulses in animals with different resections. g) Representative fluorescence trace used to calculate contraction duration. h) Duration of spontaneous contractions in animals with different resections. i) Duration of stimulated contractions in animals with different resections. N= 3 *Hydra* (GcaMP7b, Ectodermal epitheliomuscles) per resection. (Kruskal-Wallis test with dunn-sidak correction, n.s. = not significant, ** = $p < 0.01$, *** = $p < 0.001$, **** = $p < 0.0001$,***** = $p < 0.00001$,****** = $p < 0.000001$)

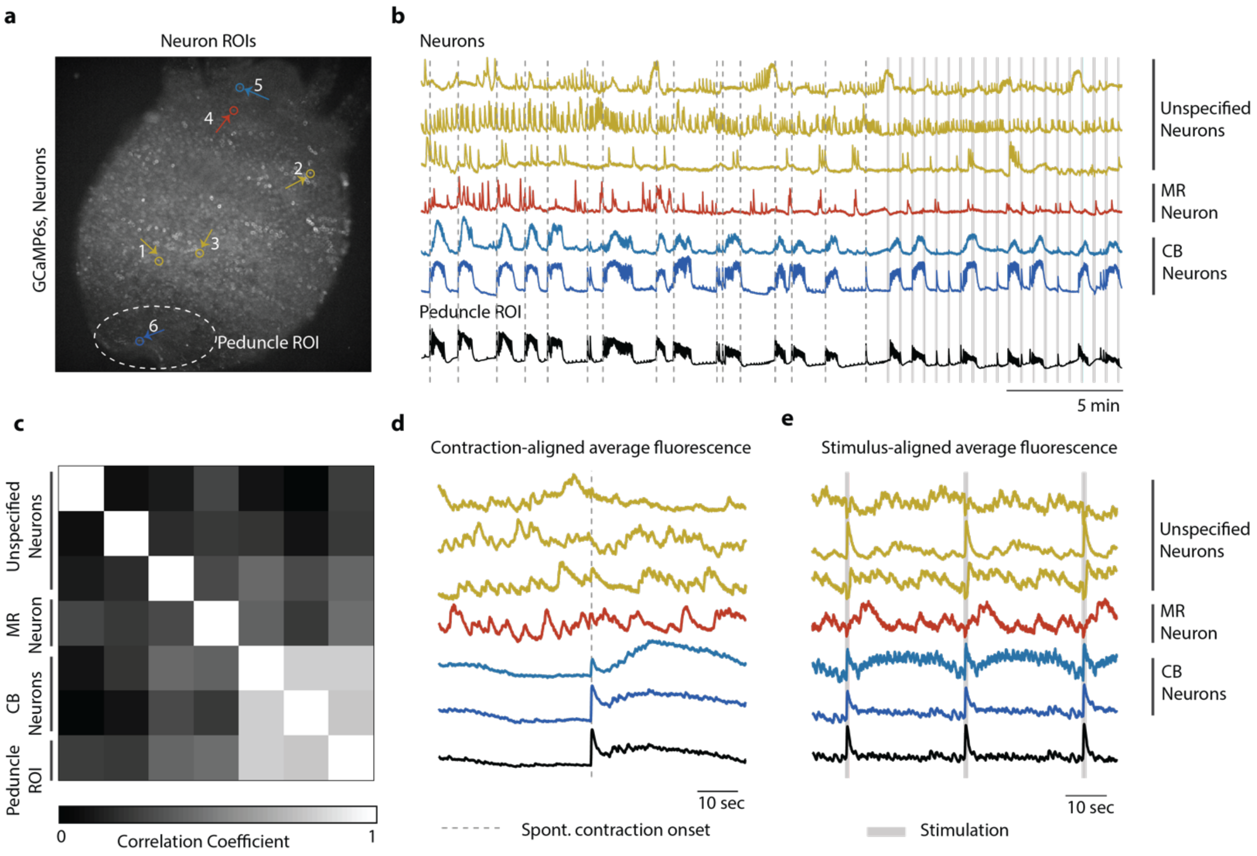


**Figure 4 – Figure Supplement 1: Single cell correlation analysis.** a) Fluorescence image of transgenic *Hydra* (#2) expressing GCaMP6s pan-neuronally. Individually tracked neuronal ROIs are indicated with circles. The neurons are numbered in the order their traces are appear in c-e with the same colors. Peduncle ROI outlined with white dashed line. b) Calcium fluorescence traces from single neurons (top 6 traces) and average calcium fluorescence from peduncle ROI (bottom trace). Mechanically responsive (MR) neurons are shown in shades of red. Contraction burst (CB) neurons are shown in shades of blue. Unspecified neurons are shown in yellow, and their activity do not resemble any of the previously identified neuronal networks (contraction burst, rhythmic potential or the mechanically responsive reported here). c) Heat map shows the correlation coefficients of individually tracked neurons and peduncle ROI. Colorbar is at the bottom. d) Average calcium fluorescence traces from each of the neurons and peduncle ROI during spontaneous behaviors time-aligned with the onset of spontaneous body contractions. Dashed line indicates the onset of body contraction. e) Average calcium fluorescence traces from each of the neurons and peduncle ROI during stimulated behaviors time-aligned with the onset of mechanical stimulation. Gray shaded rectangle indicates mechanical stimulation.

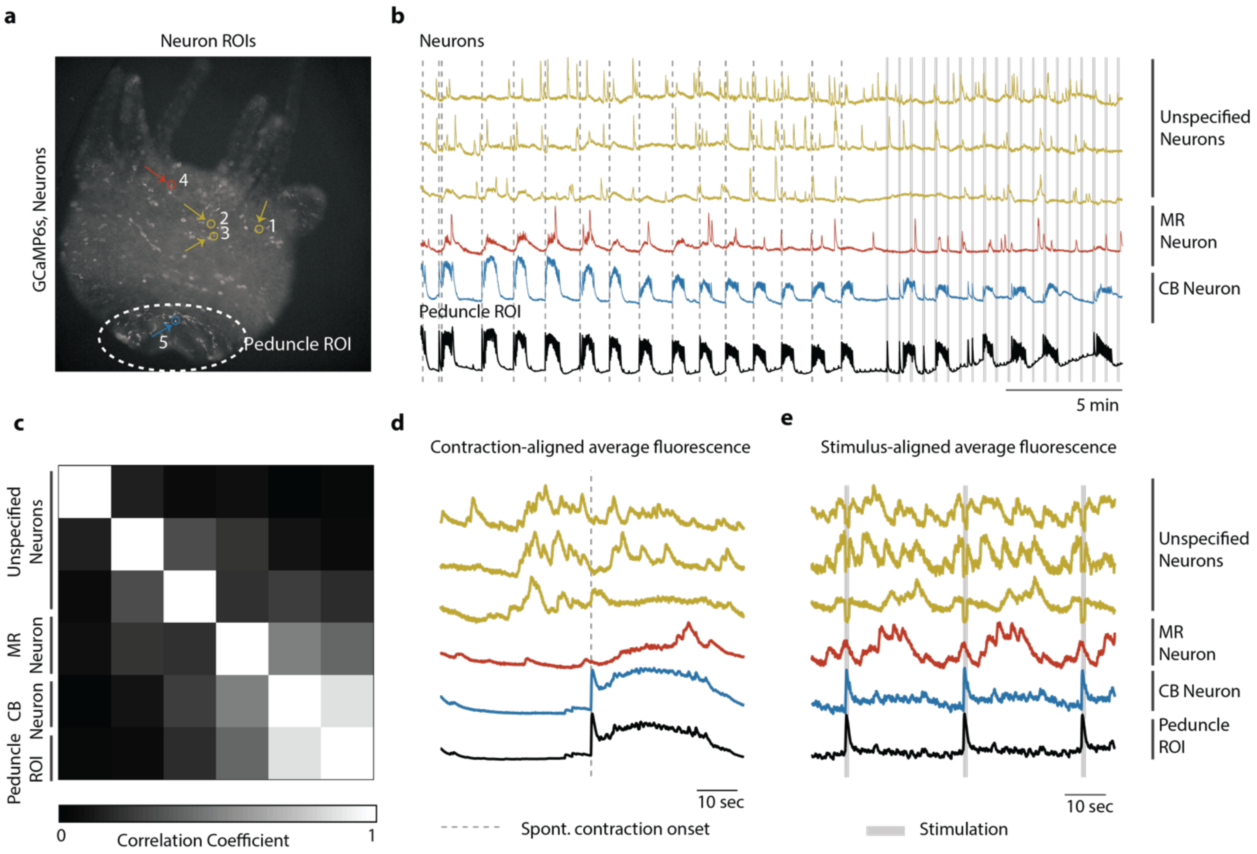


**Figure 4 – Figure Supplement 2: Single cell correlation analysis.** a) Fluorescence image of transgenic *Hydra* (#3) expressing GCaMP6s pan-neuronally. Individually tracked neuronal ROIs are indicated with circles. The neurons are numbered in the order their traces are appear in c-e with the same colors. Peduncle ROI outlined with white dashed line. b) Calcium fluorescence traces from single neurons (top 5 traces) and average calcium fluorescence from peduncle ROI (bottom trace). Mechanically responsive (MR) neurons are shown in shades of red. Contraction burst (CB) neurons are shown in shades of blue. Unspecified neurons are shown in yellow, and their activity do not resemble any of the previously identified neuronal networks (contraction burst, rhythmic potential or the mechanically responsive reported here). c) Heat map shows the correlation coefficients of individually tracked neurons and peduncle ROI. Colorbar is at the bottom. d) Average calcium fluorescence traces from each of the neurons and peduncle ROI during spontaneous behaviors time-aligned with the onset of spontaneous body contractions. Dashed line indicates the onset of body contraction. e) Average calcium fluorescence traces from each of the neurons and peduncle ROI during stimulated behaviors time-aligned with the onset of mechanical stimulation. Gray shaded rectangle indicates mechanical stimulation.

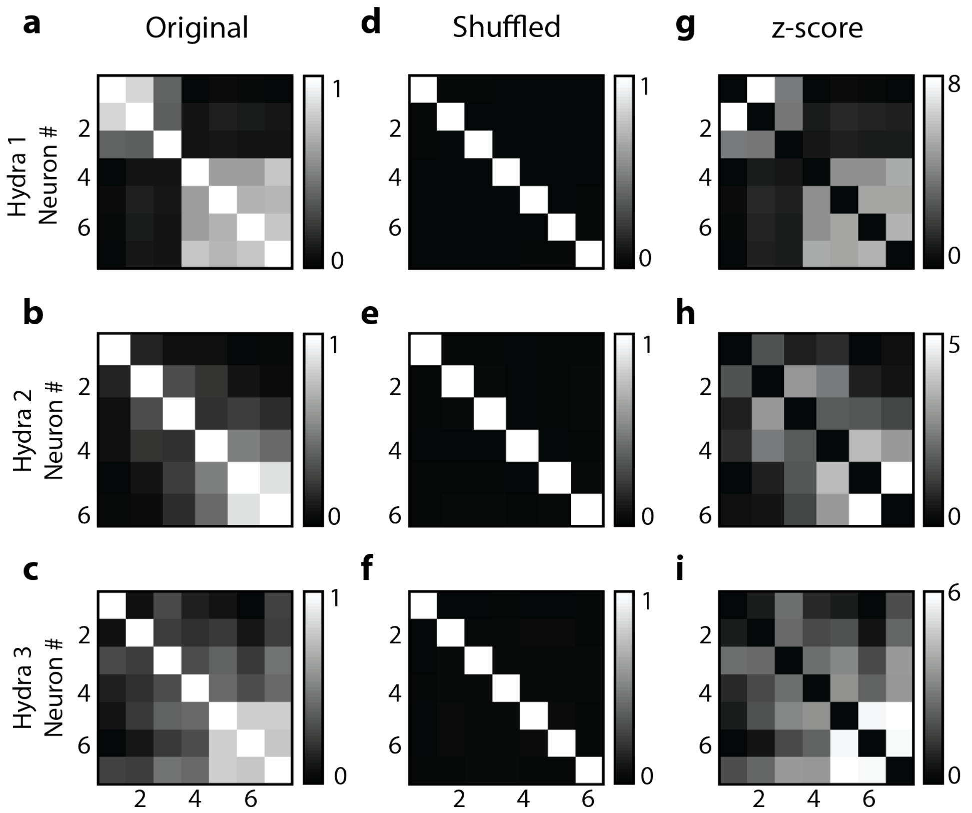


**Figure 4 – Figure Supplement 3: Random shuffling of single cell correlation analysis.** Individual neuronal fluorescence time-series were randomly shuffled to show the correlation coefficients were not due to random chance in all three *Hydra* (each row is a different *Hydra*). Original correlation coefficients from 10 mins of raw fluorescence time-series (a-c), shuffled correlation coefficients from randomly reshuffled blocks reconstructing 10 mins of fluorescence time-series (d-e) and z-score of correlation coefficients calculated based on random reshuffling (g-i). Original correlation coefficients were calculated from the 10 mins of raw fluorescence activity for each of the neurons during mechanical stimulation. Shuffled correlation coefficients were calculated by taking the mean of correlation coefficients from randomly reshuffled fluorescence time-series. Briefly, the raw fluorescence time-series from 10 mins of mechanical stimulation was divided into 100 blocks, which were then randomly recombined for each of the neurons to calculated correlation coefficients and this random reshuffling was repeated 1000 times. Z-score was calculated using the mean and standard deviation of the correlation coefficients calculated from random reshuffling repeated 1000 times. The heatmap for z-score (g-i) resemble the original correlation coefficients (a-c) indicating the correlation coefficients of the raw fluorescence time-series were not a result of random chance. Except for the diagonals which represents autocorrelation, the mean correlation coefficients resulting from randomly reshuffled fluorescence (e-f) are zero for all neuron pairs indicating the correlation due to random chance.

**Description of supplementary movies**

**Supplementary Movie 1:**

Microfluidic system to study mechanosensory response in *Hydra*. Dashed blue circle indicates the microfluidic valve that presses down on *Hydra.* Mechanosensory response from neurons and epitheliomuscular cells is shown.

**Supplementary Movie 2:**

Spontaneous neural calcium activity in normal animals. Dashed blue circle indicates the ROI used for calcium trace shown in blue (bottom). (Playback 100x.)

**Supplementary Movie 3:**

Stimulated neural calcium activity in normal animals. Dashed blue square indicates the ROI used for calcium trace shown in blue (bottom). Dashed white circle indicates the location of the valve that presses down on *Hydra* when inflated. Gray trace shows the stimulus protocol, where vertical lines indicate valve 'on' times. Stimulus applied beginning t = ~ 20 min and ends t = ~80 min. Valve is 'on' for 1 sec and 'off' for 30 sec. (Playback 100x.)

**Supplementary Movie 4:**

Spontaneous ectodermal epitheliomuscular calcium activity in normal animals. Dashed blue square indicates the ROI (entire frame) used for the calcium trace shown in blue (bottom). (Playback 100x.)

**Supplementary Movie 5:**

Stimulated ectodermal epitheliomuscular calcium activity in normal animals. Dashed blue square indicates the ROI (entire frame) used for calcium trace shown in blue (bottom). Dashed white circle indicates the location of the valve that presses down on *Hydra* when inflated. Gray trace shows the stimulus protocol, where vertical lines indicate valve 'on' times. Stimulus applied beginning t = ~ 20 min and ends t = ~80 min. Valve is 'on' for 1 sec and 'off' for 30 sec. (Playback 100x.)

**Supplementary Movie 6:**

Spontaneous neural calcium activity in longitudinally bisected animals. Dashed blue circle indicates the ROI used for calcium trace shown in blue (bottom). (Playback 100x.)

**Supplementary Movie 7:**
